# Excitatory and Inhibitory Networks Diverge Following Early Blindness

**DOI:** 10.64898/2026.02.06.704389

**Authors:** Guillaume Laliberté, Denis Boire

## Abstract

Early visual deprivation profoundly reshapes cortical functional organization, yet the contribution of distinct neuronal populations to large-scale network plasticity remains unclear. We combined awake wide-field mesoscale calcium imaging within promoter-defined neuronal populations to characterize resting-state functional connectivity in pan-neuronal (hSyn), excitatory (Thy1), and inhibitory (mDLX) cortical networks in sighted and neonatal enucleated mice. Graph-theoretical analyses revealed a convergent reorganization pattern across populations in which medial higher visual and associative cortices strengthened their connectivity with somatosensory and motor regions, whereas primary visual cortex and lateral higher visual areas lost network influence. Despite this shared motif, network remodeling differed according to neuronal identity. Excitatory networks exhibited pronounced redistribution of nodal influence and modular organization with selective alterations of global network topology, indicating selective susceptibility to sensory deprivation. Inhibitory networks preserved global efficiency while showing localized reorganization of connector and bridging hubs. Pan-neuronal networks displayed extensive redistribution of connectivity and hub architecture despite relatively preserved global network organization. These findings demonstrate that early blindness induces coordinated yet neuronal identity-dependent mesoscale network plasticity, linking mouse cortical dynamics with systems-level evidence of cross-modal reorganization in blind individuals.

## Introduction

Spontaneous brain activity plays a critical role in maintaining functional connectivity across cortical regions during rest. This intrinsic activity is organized into resting-state networks (RSNs), which reflect functional relationships shaped by underlying anatomical connectivity (Vincent et al. 2007; Greicius et al. 2009; Honey et al. 2009; Gozzi and Schwarz 2016; Whitesell et al. 2021). RSNs substantially overlap with task-evoked functional architectures and correlate with perceptual and cognitive performance, suggesting continuity between spontaneous and task-related brain organization (Lewis et al. 2009; Zhu et al. 2011; Deco et al. 2013; Kim et al. 2020). Experience and perceptual learning can further reshape these networks, indicating that spontaneous activity reflects internal representations shaped by memory and cognition.

Recent research has revealed that RSNs are not static but dynamically fluctuate over time (Oschmann and Gawryluk 2020; Özalay et al. 2024), exhibiting distinct connectivity states that correspond to changes in arousal and internal brain dynamics (Allen et al. 2014; Foster et al. 2016). This dynamic organization is further shaped by brain state and neuronal identity-dependent properties, particularly through the distinct contributions of excitatory and inhibitory neuronal populations to large-scale functional brain architecture, as demonstrated in animal models (O’Connor et al. 2022).

Importantly, this intrinsic functional architecture is sensitive to sensory experience. Alterations in functional networks among individuals deprived of sensory input early in life, including those with early blindness, provide compelling evidence that sensory input is required for typical network development. Studies in both humans and animals have shown that sensory loss can lead to large-scale reconfiguration of resting-state networks, reflecting cross-modal plasticity (Liu et al. 2007; Yu et al. 2008; Burton et al. 2014; Hou et al. 2017; Kraft et al. 2017; Pelland et al. 2017; Shao et al. 2018; Huang et al. 2019; Li et al. 2019; Guerreiro et al. 2021; Wang et al. 2022; Chen et al. 2024; Laliberté and Boire 2026) leading to the recruitment of visual cortical areas by non-visual modalities (Bedny et al. 2011; Ptito et al. 2012; Kanjlia et al. 2016; van der Heijden et al. 2020; Kanjlia et al. 2021) and improved discrimination capabilities in the conserved modalities (Amedi et al. 2004; Collignon et al. 2009; Lazzouni et al. 2012).

These observations support the hypothesis of a metamodal cortex dependent at least in part on optimal sensory input for its development (Pascual-Leone and Hamilton 2001; Bedny 2017; Kanjlia et al. 2019; Saccone et al. 2024). This reorganization does not appear to rely on the emergence of new anatomical connections, but rather on the selective potentiation and unmasking of pre-existing multisensory and intracortical pathways through Hebbian and homeostatic mechanisms (Lee and Whitt 2015; Ewall et al. 2021; Makin and Krakauer 2023), notably descending multisensory projections and latent intracortical pathways (Meng et al. 2015; Petrus et al. 2015). In line with this view, Makin and Krakauer (2023) propose that cross-modal plasticity primarily reflects the amplification of latent computational capacities embedded within existing cortical architectures rather than extensive structural rewiring of cortical circuitry.

During critical periods of postnatal development, synaptic remodeling in sensory cortices is regulated by the maturation of GABAergic interneurons and the establishment of excitation/inhibition (E/I) balance (Hensch et al. 1998; Fagiolini and Hensch 2000; Hensch 2005; Sale et al. 2010). Inhibitory microcircuits refine sensory responses locally and regulate feedforward excitation within cortical networks (Tao and Poo 2005; Iurilli et al. 2012; Li et al. 2014; Wu et al. 2022). Visual experience is known to shape E/I dynamics not only in visual cortex (Benevento et al. 1992; Morales et al. 2002; Chattopadhyaya et al. 2004; Hensch and Fagiolini 2005; Gandhi et al. 2008; Desgent et al. 2010; Kameyama et al. 2010; Levelt and Hubener 2012; Kannan et al. 2016; Cheng et al. 2022) but also in non-visual regions (Goel et al. 2006; Carriere et al. 2007; Mowery et al. 2016; Nakajima et al. 2016). Consistent with this framework, visual deprivation alters inhibitory and excitatory signaling in the visual cortex, including shifts in Glx/GABA ratio, reduced inhibitory tone, and enhanced corticocortical excitation (Goel *et al*. 2006; Jiao et al. 2006; Cheetham et al. 2007; Petrus *et al*. 2015; Rączy et al. 2022; Pant et al. 2025). These changes are thought to promote heightened plasticity and increased sensitivity to non-visual inputs within deprived visual regions.

Following visual deprivation, preserved sensory modalities exhibit enhanced local inhibition, which sharpens unimodal processing and limits cross-modal interference (Petrus et al. 2014). In parallel, the deprived visual cortex undergoes enhanced corticocortical excitation and reduced inhibitory control, resulting in a shift in E/I balance that may promote heightened plasticity and increased sensitivity to non-visual inputs (Goel *et al*. 2006; Petrus *et al*. 2015; Lee 2022). Consistent with these observations, both animal and human studies report increased excitability and functional recruitment of deprived visual cortices following early blindness (Röder et al. 2002; Weaver et al. 2013; Coullon et al. 2015; Bedny 2017; Rączy *et al*. 2022; Pant *et al*. 2025).

Together, inhibitory-dominant adaptations in spared modalities (Meng *et al*. 2015, 2017) and excitatory-driven remodeling in the deprived cortex suggest that sensory deprivation induces asymmetric E/I reorganization across cortical networks. Enhanced excitation and reduced inhibition within deprived visual regions may facilitate integrative plasticity and cross-modal recruitment, whereas preserved sensory cortices undergo inhibitory reinforcement that may support modality-specific processing and limit interference from competing inputs. These asymmetric E/I adjustments may therefore reflect large-scale functional reorganization mediated through the potentiation and redistribution of pre-existing cortical pathways rather than the formation of new anatomical circuitry.

Despite this wealth of evidence, the neuronal identity-dependent mechanisms behind RSN reorganization in sensory-deprived individuals remain poorly understood. In particular, it is unclear how excitatory and inhibitory cortical neuronal subpopulations each contribute to large-scale network remodeling that follows early vision loss, and whether their differential engagement underlies shifts in integration, segregation, and hub architecture.

In this study, we address this issue with wide-field mesoscale calcium imaging targeted to pan-neuronal, glutamatergic, or GABAergic neuronal populations through specific neuronal promoters (hSyn, Thy1 and mDLX) following neonatal enucleation in mice. We quantified resting-state connectivity, graph-theoretic metrics, and modular organization across these neuronal populations. We hypothesized that early visual deprivation would induce neuronal identity-dependent reorganization of cortical networks. Specifically, we expected excitatory, inhibitory, and pan-neuronal populations to exhibit distinct mesoscale connectivity patterns following loss of visual input, reflecting their unique roles in shaping network integration and modularity during critical developmental windows.

## Materials and Methods

### Mice

All experimental procedures were approved by the Animal Care Committee of the Université du Québec à Trois-Rivières (CBSA, protocol DB17) and adhered to the guidelines of the Canadian Council on Animal Care. A total of 33 C57BL/6J-Tg (Thy1-GCaMP6s)GP4.3Dkim/J mice (JAX stock #024275) were used. Of these, 12 mice carried the Thy1-GCaMP6s transgene, enabling the expression of the calcium reporter selectively in projecting excitatory neurons (Porrero et al. 2010; Dana et al. 2014), and 21 were non-transgenic littermates.

Mice were divided into six experimental groups according to calcium indicator expression and visual condition: sighted C57BL/6J hSyn-GCaMP6s mice (n = 6; 2 females/4 males), enucleated C57BL/6J hSyn-GCaMP6s mice (n = 3; 1 female/2 males), sighted C57BL/6J Thy1-GCaMP6s mice (n = 6; 2 females/4 males), enucleated C57BL/6J Thy1-GCaMP6s mice (n = 6; 2 females/4 males), sighted C57BL/6J mDLX-GCaMP6s mice (n = 5; 2 females/3 males), and enucleated C57BL/6J mDLX-GCaMP6s mice (n = 7; 3 females/4 males). Sex distribution was balanced as much as possible across experimental groups; however, the study was not statistically powered to assess sex-dependent effects. The complete experimental timeline is shown in **Figure 1A**. Mice were weaned at three weeks of age, and, except for neonatal enucleation, experimental procedures began at six weeks of age.

**Figure 1.**
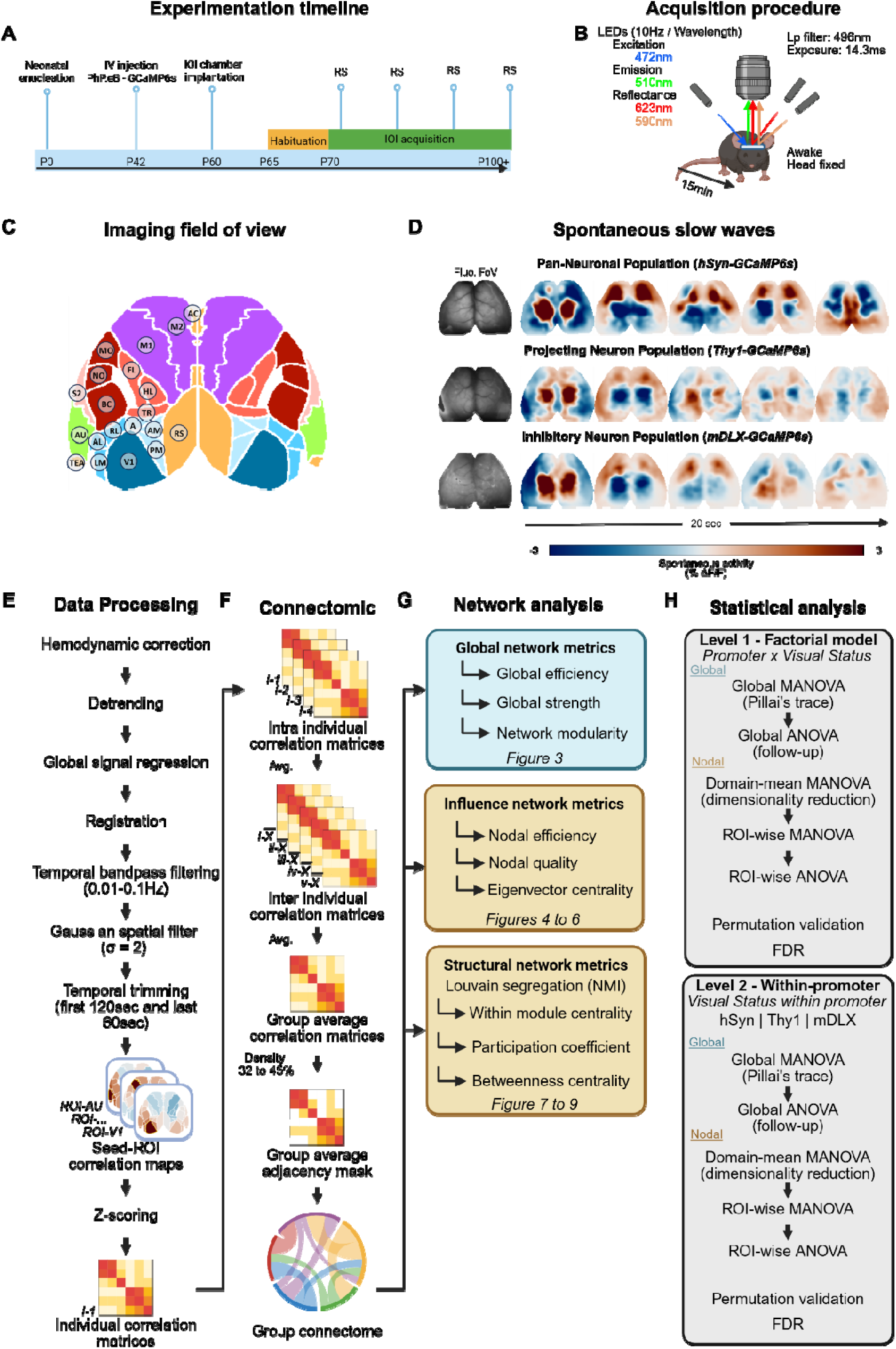
– **Experimental timeline, acquisition workflow, preprocessing pipeline, and graph-theoretical analysis of wide-field calcium imaging in awake mice. A.** Experimental timeline showing neonatal enucleation, viral delivery, chronic imaging chamber implantation, habituation, and repeated resting state (RS) recordings between P70 and P90. B. Wide-field mesoscale calcium imaging setup using sequential illumination for fluorescence excitation and hemodynamic correction. C. Cortical parcellation adapted from the Allen Brain Atlas. Functional domains are color-coded as motor (purple), associative (yellow), somatosensory (red), visual (blue), and auditory (green). D. Representative spontaneous cortical activity patterns across pan-neuronal (hSyn-GCaMP6s), excitatory (Thy1-GCaMP6s), and inhibitory (mDLX-GCaMP6s) populations. E. Preprocessing workflow including hemodynamic correction, filtering, registration, global signal regression, and ROI-based correlation analysis. F. Generation of functional connectivity matrices and adjacency masks for connectomic analyses. G. Graph-theoretical analysis pipeline illustrating global, nodal influence, and community metrics. H. Statistical analysis workflow including MANOVA, permutation testing, and FDR correction *(Created in BioRender. LALIBERTÉ, G. (2026)* https://BioRender.com/jfmfjcb).

Data collection occurred between three and six months of age. Animals were maintained under a 12-hour light/dark cycle with ad libitum access to food and water. Inclusion and exclusion criteria were established a priori to ensure data quality and optical stability. Only animals exhibiting consistent and robust mesoscale fluorescence signals across all imaging sessions were included. For sighted controls, ocular integrity had to remain intact throughout the entire experimental timeline, with no evidence of corneal opacity or surface alteration. Furthermore, only datasets from animals that completed the full imaging protocol were analyzed, which required maintaining a clean, stable, and artifact-free imaging chamber without the development of optical distortions. In this study, no animals met exclusion criteria, and all thirty-three mice were retained for analysis.

### Enucleation

At parturition, dams received a subcutaneous injection of long-acting Buprenorphine SR-LAB (1.0 mg/kg; Chiron Compounding Pharmacy Inc., Guelph, ON). Within 24 hours postpartum, bilateral enucleation was performed on half of the pups under deep hypothermia-induced anesthesia. The palpebral fissure was carefully opened with a scalpel, and the optic nerve was severed to extract the eyeball. The ocular cavity was filled with hemostatic gel foam (Spongostan; Johnson & Johnson, New Brunswick, NJ, USA), and the eyelids were sealed with tissue adhesive (Vetbond; 3M, St. Paul, MN, USA). Pups were rewarmed until full recovery and returned to their home cage.

### Intravenous Viral Vector Injection

To enable GCaMP6s expression in non-transgenic mice, adeno-associated viral vectors (AAV2/PHP.eB capsids) were used to deliver plasmids encoding the calcium reporter. Two promoter systems were employed: AAV-hSyn-GCaMP6s (construct-889-aavphp-eb), enabling broad neuronal expression under the human synapsin promoter, or AAV-mDLX-GCaMP6s (construct-880-aavphp-eb), driving expression in GABAergic interneurons under the mDLX enhancer. The synapsin promoter confers pan-neuronal expression in the cortex while avoiding glial transduction (Kugler et al. 2003; Jin et al. 2016; Finneran et al. 2021). The mDLX enhancer ensures forebrain-specific expression in inhibitory interneurons (Dimidschstein et al. 2016). Moreover, mDLX allows targeting interneurons broadly and uniformly as PV, SST and VIP GABAergic neuronal populations can be labeled in proportions corresponding to these three populations in cortex (Dimidschstein *et al*. 2016).

Viral vectors were provided by the Canadian Neurophotonics Platform Viral Vector Core Facility (RRID:SCR_016477) and injected intravenously into the caudal vein of 42-day-old C57BL/6J mice. Each injection consisted of 300 μL of sterile phosphate-buffered saline (PBS) containing 6.0 × 10¹¹ genome copies. The AAV-PHP.eB serotype facilitates efficient, widespread neuronal transduction across the cortex by enhancing blood-brain barrier penetration (Chan et al. 2017; Matsuzaki et al. 2019; Michelson et al. 2019).

Because neonatal enucleation was performed at birth and imaging acquisitions began several weeks later (P70–P90), acute inflammatory responses associated with optic nerve injury were expected to have resolved before recordings. Similarly, AAV vectors were administered intravenously rather than through direct cortical injections, reducing the likelihood of focal cortical inflammatory responses.

### Implantation of Chronic Imaging Chambers

Chronic imaging chambers were installed when mice were at P60. Mice received a subcutaneous injection of long-acting Buprenorphine SR-LAB (1.0 mg/kg), followed by anesthesia induction with isoflurane (5% induction, 2% maintenance in medical grade O ). Animals were positioned in a stereotaxic frame, and body temperature was maintained at 37°C with a heating pad and monitored with a rectal thermometer. Eyes were protected from corneal dehydration using veterinary ophthalmic gel (Aventix Pet Eye Lube, Aventix Animal Health, Burlington, ON, Canada), as needed.

The scalp was shaved and disinfected using sequential application of 2% chlorhexidine, 70% ethanol, and 16% iodine. Local anesthesia was provided by subcutaneous injections of Lidocaine 2% (7□mg/kg) combined with Bupivacaine 0.25% (3.5□mg/kg). The skull was covered with transparent dental cement (C&B MetaBond; Parkell, Edgewood, NY, USA), sealed with a glass coverslip (Carolina; Burlington, NC, USA), and affixed with a titanium head bar (1.1□g, 25□×□3.2□×□3.2□mm; SKU: 515301; Labeo Technologies Inc.).

Following surgery, animals received subcutaneous carprofen (0.5 mg/kg) and were housed individually for recovery. Additional doses of carprofen (0.5 mg/kg) were administered at 24-, 48-, and 72-hour post-surgery. After 5 days, females were housed in groups of 3–5, and males were paired in cages separated by clear plexiglass dividers to promote social interaction while preventing aggression.

### Mesoscale Calcium Imaging Recordings

To minimize circadian variability, all imaging sessions were conducted between 8:00 AM and 2:00 PM. Mice were habituated to the head-fixation apparatus over five consecutive days (P60 to P65): Day 1 involved 5 minutes on the setup without head fixation; Days 2–5 introduced increasing duration of the head fixation (5, 10, 20, and 40 minutes). Brain illumination remained off for the first four days and was activated during the final session (**Figure 1A**). This habituation protocol was designed to minimize behavioral stress during subsequent imaging sessions. Recordings, starting at P70, were performed under a wide-field mesoscale imaging system (OiS200; Labeo Technologies) in complete darkness. Calcium signals were captured using a vertically mounted CCD camera equipped with NIKKOR 50 mm f/1.2 lenses (Nikon, Minato, Tokyo, Japan). GCaMP6s indicators were excited with 472 nm blue light (Cree XLamp XP-E2; Cree, Durham, NC, USA), whereas the 590 nm (Amber LED, LZ4-00MA00) and 623 nm (Red LED, LZ4-00MA00; OSRAM, Markham, ON, Canada) channels were used to acquire oxyhemoglobin- and deoxyhemoglobin-related reflectance signals for subsequent hemodynamic correction (**Figure 1B**). To prevent excitation light from reaching the eyes and to minimize potential sensory confounds, an opaque black occluder was positioned around the imaging field and illumination was restricted to the boundaries of the chronic cranial window.

Image acquisition occurred at 30 Hz (10 Hz per wavelength to support hemodynamic correction) with a resolution of 512 × 512 pixels and 14.33 ms exposure time. A 496 nm long-pass filter was mounted to the objective to prevent spectral crosstalk. Each mouse underwent four 15-minute resting-state recordings in a quiet, dark environment, spaced across 4 to 6 weeks **(Figure 1A)**.

### Data Processing and Analysis of Calcium Signals

Cortical regions were delineated by aligning the top-projection mask from the Mouse Allen Brain Atlas (**Figure 1C**). Masks were scaled to the pixel/mm ratio and eroded by 5 pixels to enhance regional specificity. Twenty cortical ROIs were identified per hemisphere and classified according to their predominant functional organization: association cortex (AC, RS, TEA), motor cortex (M1, M2), dorsal column-related somatosensory cortex (HL, TR, FL), trigeminal-related somatosensory cortex (BC, NO, MO), secondary somatosensory cortex (S2), medial visual cortex (PM, AM, A), primary visual cortex (V1), lateral visual cortex (RL, AL, LM), and primary auditory cortex (AU). Representative examples of cortical infraslow waves recorded from these regions are shown in **Figure 1D**.

Data preprocessing was performed using the Universal Mesoscale Imaging Toolbox (UmIT Bipolar-v2.0.0; available at labeotech.github.io/Umit; Ferreira de Souza et al. 2025) following the workflow illustrated in **Figure 1E**. Wavelength-sorted image stacks were spatially down sampled to 128 × 128 pixels (92.2 μm/pixel) using *run_ImagesClassification*. Hemodynamic signal correction was performed using *run_HemoCorrection* according to Valley et al. (2020). Briefly, pixel-wise linear regression of the fluorescence signal against the reflectance channels was used to remove non-neuronal hemodynamic contamination arising from fluctuations in cerebral blood volume, hemoglobin oxygenation, light absorption, and scattering associated with vascular dynamics, thereby isolating neuronal calcium-dependent activity. Time series were detrended using *apply_detrend*, and global signal regression (using *GSR*) was applied across manually defined cortical surfaces to reduce non-neuronal fluctuations, including cardiac and respiratory artifacts (Turley et al. 2017). GSR was further implemented to attenuate widespread fluctuations shared across the cortical surface and improve the specificity of region-dependent functional correlations. Although GSR can modify the covariance structure of neural signals and induce negative correlations (Murphy et al. 2009; Murphy and Fox 2017), it was retained because awake mesoscale calcium imaging is particularly susceptible to global signal contamination arising from animal movement, respiration, photobleaching, LED intensity drift, and optical inhomogeneity associated with transcranial imaging (see Brier and Culver 2023). The use of GSR in this context is consistent with previous mesoscale calcium imaging studies of mouse cortex (Vanni and Murphy 2014; Vanni et al. 2017; Wright et al. 2017; Brier et al. 2019; O’Connor *et al*. 2022). Image sequences were aligned to a reference frame using *alignFrames*, and registration accuracy was manually verified. Signals were then temporally filtered between 0.0083 and 0.1 Hz using *normalizeLPF* and spatially smoothed with a Gaussian filter (σ = 2) using *spatialGaussFilt*. To minimize potential habituation effects, the first 120 s and the final 60 s of each recording session were excluded from further analyses using *trim_movie*. ROI-averaged calcium time series were subsequently extracted by averaging all pixels contained within each cortical mask for downstream functional connectivity analyses.

### Network Analysis and Rationale

Functional connectivity matrices were constructed by computing Pearson correlation coefficients between ROI time series across both hemispheres. Correlation coefficients were Fisher z-transformed using *genCorrelationMatrix* in UmIT before graph-theoretical analysis. Because the graph analyses were designed to quantify the magnitude of interregional functional coupling independently of correlation direction, the absolute values of the Fisher z-transformed coefficients were used as edge weights. Accordingly, strongly positive and strongly negative correlations were both represented as strong functional associations, whereas correlations approaching zero were represented as weak associations. Proportional thresholding was subsequently applied to these absolute weighted matrices, retaining connections according to their coupling magnitude. The resulting graphs therefore represented unsigned functional-connectivity magnitude networks and did not preserve information regarding the original direction of the correlations. Networks comprised 20 ROIs per hemisphere and were thresholded across proportional network densities ranging from 10% to 45%, in 1% increments, to derive the final adjacency matrices (**Figure 1F**).

Graph metrics were computed using the Brain Connectivity Toolbox (Rubinov and Sporns 2010), including global efficiency, nodal strength, eigenvector centrality, modularity, participation coefficient, betweenness centrality, and within-module degree z-score (**Figure 1G**). Within this unsigned framework, global and nodal strength quantified the magnitude of functional coupling, whereas global efficiency characterized network integration based on weighted paths derived from absolute connectivity magnitudes. Modularity, participation coefficient, betweenness centrality, and within-module degree z-score were used to characterize community structure and the structural roles of individual cortical regions. Together, these measures provided a multiscale characterization of global integration, nodal influence, and modular organization following early visual deprivation (Laliberté and Boire 2026).

### Statistical Analysis

All statistical analyses were performed in R (version 4.6.0; R Foundation for Statistical Computing, Vienna, Austria) using RStudio (version 2026.04.0; Posit Software, Boston, MA, USA), according to the statistical pipeline illustrated in **Figure 1H**.

The mouse served as the experimental unit. Connectivity matrices and graph-theoretical metrics were averaged across recording sessions within each animal prior to group-level analyses to avoid pseudoreplication. Hemispheric comparability was first assessed using permutation testing on correlation matrices; because no significant hemispheric differences were detected, metrics were averaged across hemispheres for each animal.

Graph metrics were computed across density thresholds (32–45%) and averaged across thresholds to reduce dependence on arbitrary threshold selection. The lower bound corresponded to the minimum density at which all nodes remained connected, whereas the upper bound avoided excessively dense networks approaching fully connected graphs (Fornito et al. 2016). Nodes were classified as hubs when their metric values exceeded the ROI mean, ranked within the upper 15% of nodes, and remained consistently identified across at least 70% of density thresholds.

Metrics were grouped into global, nodal influence, and nodal structural categories. Global effects were assessed using factorial MANOVA (Pillai’s trace), testing visual status, promoter population, and their interaction, followed by univariate ANOVAs for significant multivariate effects. To reduce dimensionality relative to sample size, nodal analyses employed a hierarchical framework in which metrics were first averaged within predefined functional domains prior to domain-level MANOVAs. Domains showing significant effects after multiple-comparison correction were subsequently examined at the ROI level. ROI-level inferential results are therefore reported only for domains that passed this hierarchical domain-level criterion. Regional patterns from domains that did not meet this criterion, including changes in hub classification, are presented descriptively and were not subjected to independent statistical inference.

Permutation testing was used to validate MANOVAs and ANOVAs. For within-population effects, visual-status labels were permuted while preserving group sizes; for interaction effects, permutations were restricted within promoter groups. Effect sizes were estimated using partial eta squared, and statistical significance was defined as FDR-corrected p ≤ 0.05 (Benjamini– Hochberg procedure).

Figures were generated using custom MATLAB scripts (version 2024.2; The MathWorks, Inc.). Representative connectomes and community structures are shown at 38% network density, whereas all statistical analyses were performed on metrics averaged across densities (32–45%).

## Results

In this study, we investigated spontaneous resting-state cortical activity in awake mice expressing the calcium indicator GCaMP6s across three neuronal populations: pan-neuronal (hSyn promoter), excitatory neurons (Thy1 promoter), and inhibitory neurons (mDLX mini promoter).

## Baseline Event Dynamics Across Neuronal Populations

To assess cross-cohort comparability and potential differences related to promoter systems and expression strategies, spontaneous calcium event dynamics were quantified by detecting positive and negative events from z-score-normalized calcium traces (±1 SD threshold, minimum inter-event interval = 0.3 s). Event frequency and mean peak amplitude were compared across promoter groups (**Supplementary Figure 1**).

Inter-promoter comparisons within sighted animals revealed no significant differences across cortical domains after FDR correction. These findings indicate that spontaneous event dynamics remained broadly comparable across promoter systems. Visual deprivation induced population-dependent modulation of activity dynamics within visual cortical regions.

## Alterations in Functional Connectivity Patterns

To provide an exploratory overview of connectivity changes induced by early visual deprivation, we compared intra- and interhemispheric resting-state functional connectivity between sighted and enucleated mice using uncorrected Welch’s *t*-tests (p < 0.05). This analysis revealed widespread but heterogeneous increases and decreases in functional connectivity across the cortex, suggesting extensive reorganization of both local and long-range networks (**Figure 2**).

**Figure 2.**
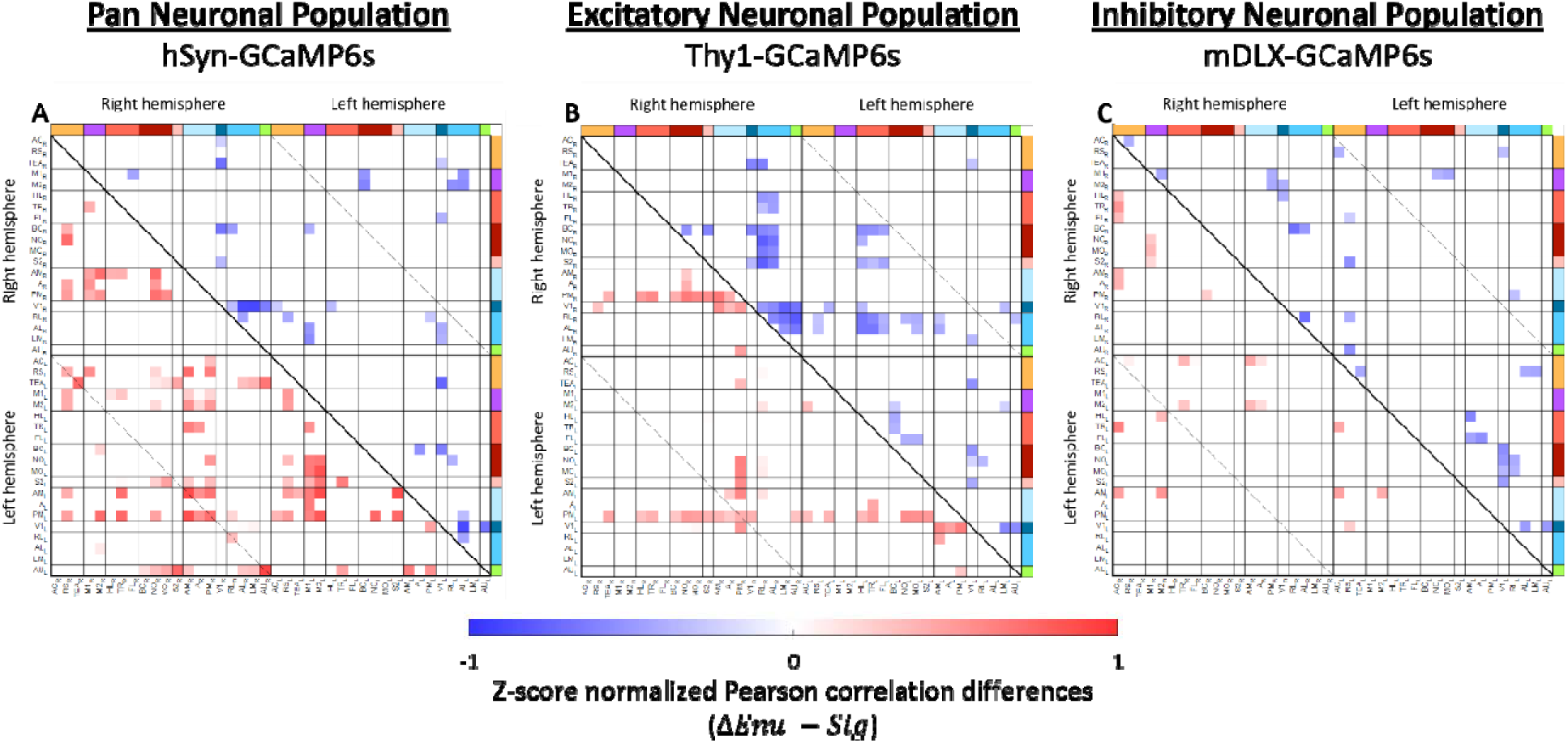
– Altered Functional Connectivity Following Early Visual Deprivation. Significant inter-regional correlation differences (Welch’s t-test, p < 0.05, uncorrected) between early visually deprived and sighted mice. Positive differences (red shades) are shown in the lower triangle, negative differences (blue shades) in the upper triangle. Panels depict: **A.** pan-neuronal population (hSyn-GCaMP6s), **B.** projecting (excitatory) neuronal population (Thy1-GCaMP6s), and **C.** inhibitory neuronal population (mDLX-GCaMP6s), comparing sighted and enucleated mice. Brain regions are organized by hemisphere (right, left) and by functional domains—associative, motor, somatosensory, visual, and auditory, color-coded as described in Figure 1C.

### Pan-Neuronal Population (hSyn-GCaMP6s)

Early visual deprivation induced widespread reorganization of pan-neuronal functional connectivity (**Figure 2A**). Interhemispheric connectivity was predominantly strengthened, particularly between medial visual, associative, motor, and somatosensory regions, whereas reductions were mainly restricted to motor-sensory interactions and connections involving V1 and lateral visual cortices. Similarly, intrahemispheric connectivity showed increased coupling between medial higher-order visual areas and associative, motor, and somatosensory cortices, while connectivity losses were largely confined to V1 and lateral visual networks.

### Projecting (Excitatory) Neuronal Population (Thy1-GCaMP6s)

Within Thy1 networks, early visual deprivation was associated with a spatially selective pattern of functional connectivity remodeling (**Figure 2B**). Interhemispheric increases were largely driven by the medial visual area PM, whereas connectivity reductions remained limited. Similarly, intrahemispheric connectivity was characterized by strengthened coupling between medial visual areas and multiple cortical systems, while V1 exhibited reduced connectivity primarily with lateral visual and selected associative and somatosensory regions.

### Inhibitory Neuronal Population (mDLX-GCaMP6s)

Within inhibitory (mDLX) networks, functional connectivity remained largely preserved following early visual deprivation, with spatially restricted changes (**Figure 2C**). Interhemispheric connectivity showed only limited changes, with modest increases involving associative and motor cortices and few localized decreases. Likewise, intrahemispheric connectivity exhibited selective strengthening between associative, medial visual, and somatosensory regions, whereas connectivity reductions were largely restricted to interactions involving visual and sensorimotor cortices.

## Global and Cell-Identity Dependent Network Reorganization Following Early Visual Deprivation

To determine whether early visual deprivation differentially reorganizes large-scale cortical network topology across neuronal populations, graph-theoretical analyses were performed on weighted functional connectomes across proportional density thresholds (32–45%). Representative difference connectomes are shown at 38% network density, corresponding to the midpoint of the analyzed range (**Figure 3A).**

**Figure 3.**
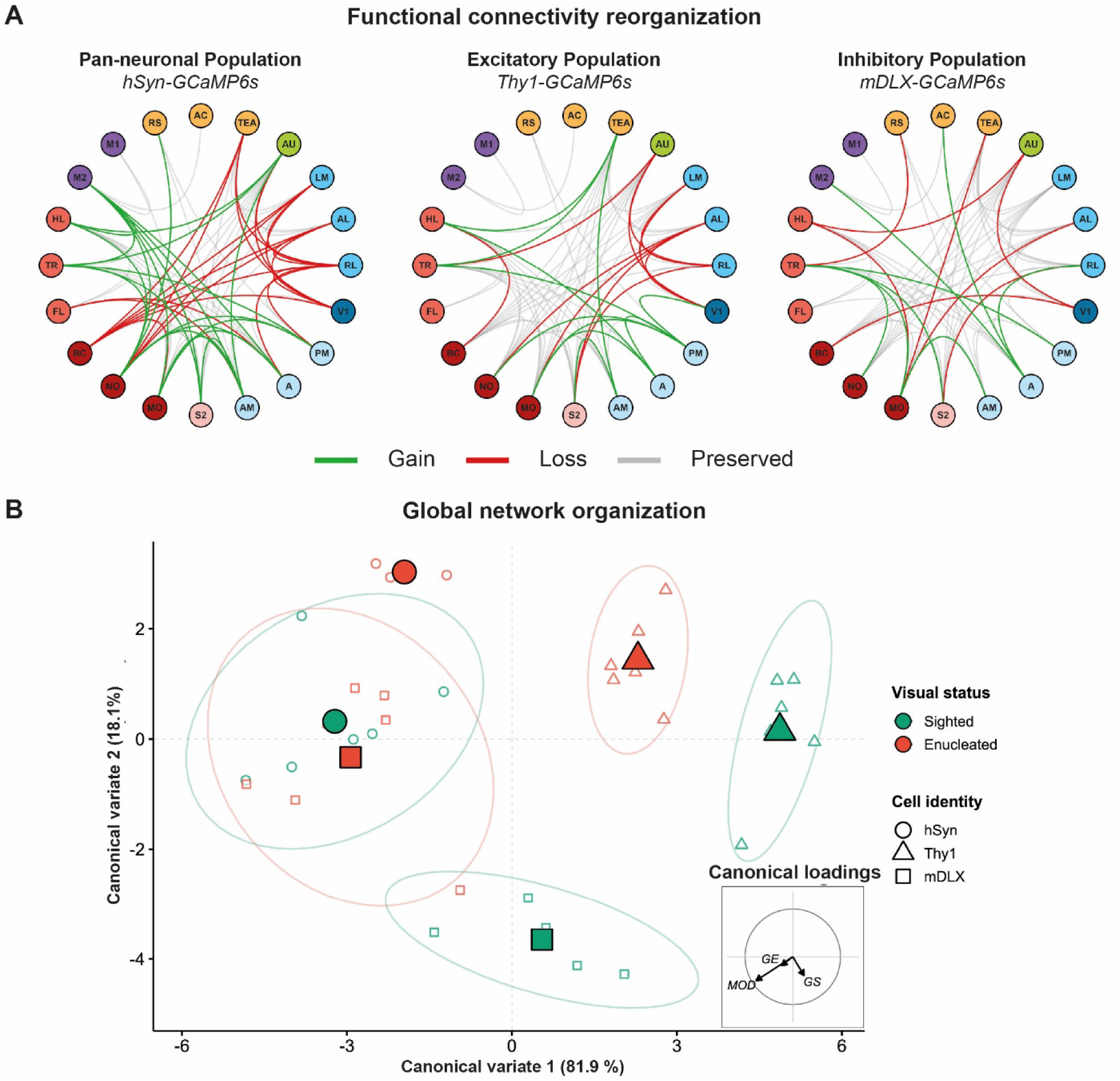
– Functional connectivity reorganization and global network organization following early visual deprivation. Effects of neonatal enucleation on pan-neuronal (hSyn-GCaMP6s), excitatory (Thy1-GCaMP6s), and inhibitory (mDLX-GCaMP6s) cortical functional networks in C57BL/6J mice. **A.** Functional connectivity reorganization. Circular connectome diagrams illustrate suprathreshold functional connectivity changes (network density = 38%) between cortical regions. Connections are classified as gained, lost, or preserved in early blind compared to sighted mice. Nodes are grouped according to the functional cortical modules defined in Figure 1C. **B. Global network organization.** Canonical discriminant analysis of standardized global efficiency, Louvain modularity, and global connectivity strength, applied to the Visual Status × Neuronal Population interaction term from the factorial MANOVA. Each small symbol represents the canonical scores of one animal. Symbol shape identifies the neuronal population, whereas color indicates visual status. Large, filled symbols represent group centroids, and normal-theory ellipses encompass 80% of the observations within each Visual Status × Neuronal Population group. The inset correlation circle displays the structure correlations between the standardized global network metrics and the first two canonical variates (GE, global efficiency; GS, global strength; MOD, Louvain modularity). Because the enucleated hSyn group included only three animals, a normal-theory covariance ellipse could not be estimated reliably and is therefore not displayed for this group.

Global network organization was assessed using global efficiency, modularity, and total connectivity strength, reflecting integration (Rubinov and Sporns 2010), segregation (Blondel et al. 2008) and overall coupling (van den Heuvel and Sporns 2011), respectively across pan-neuronal (hSyn), excitatory (Thy1), and inhibitory (mDLX) networks (**Figure 3B**).

### Cell Identity–Dependent Global Network Reorganization Following Early Visual Deprivation

Multivariate analysis of global network metrics revealed significant effects of neuronal population (Pillai’s trace = 1.59, F(6,52) = 33.41, p < 0.001) and visual status (Pillai’s trace = 0.78, F(3,25) = 28.90, p < 0.001), together with a significant Neuronal Population × Visual Status interaction (Pillai’s trace = 0.71, F(6,52) = 4.72, p = 0.001; permutation p = 0.002).

To visualize the multivariate structure underlying the significant interaction, CDA was applied specifically to the Visual Status × Neuronal Population interaction term from the fitted factorial MANOVA (Figure 3B). The first and second canonical variates accounted for 81.9% and 18.1% of the interaction-related discriminant variation, respectively. The neuronal populations occupied distinct regions of canonical space and exhibited different deprivation-related trajectories. Visual deprivation shifted the Thy1 and mDLX centroids toward lower Can1 and higher Can2 scores, whereas the hSyn centroid shifted toward higher scores on both axes, predominantly along Can2. Structure correlations indicated that Can1 primarily contrasted modularity (r = −0.768) with global strength (r = 0.237), with a weaker contribution from global efficiency (r = −0.236). All three metrics were negatively correlated with Can2, with the strongest association observed for modularity (r = −0.494), followed by global strength (r = −0.391) and global efficiency (r = −0.181). Thus, the canonical representation illustrated population-dependent differences in the balance among network segregation, integration, and overall functional coupling, with modularity contributing most strongly to the multivariate interaction.

Follow-up MANOVAs within each neuronal population identified significant effects of visual deprivation in Thy1 (Pillai’s trace = 0.94, F(3,8) = 44.18, p < 0.001) and mDLX networks (Pillai’s trace = 0.82, F(3,8) = 12.34, p = 0.0023). A weaker effect was detected in hSyn networks (Pillai’s trace = 0.77, F(3,5) = 5.59, p = 0.047) but did not remain significant under permutation testing. Consistent with the structure correlations, univariate analysis indicated that the interaction was primarily driven by modularity (F(2,27) = 7.84, p = 0.0021; permutation p = 0.0070; η² = 0.37), whereas the effect on global connectivity strength did not survive permutation correction.

To contextualize these population-dependent global patterns, representative difference connectomes were examined for each neuronal population (**Figure 3A**). Subsequent nodal influence and modular analyses were then used to localize the regional network reorganization associated with these distinct global trajectories.

### Pan-Neuronal Population (hSyn-GCaMP6s)

In hSyn-expressing enucleated mice (**Figure 3A**), cortical connectivity underwent widespread redistribution. V1 preferentially lost connections with associative, auditory, trigeminal somatosensory, and lateral visual cortices while preserving its connectivity with medial visual areas. Similarly, lateral visual cortices exhibited extensive losses of both intra-visual and cross-modal connectivity. In contrast, medial visual areas (PM, A, and AM) gained connectivity with somatosensory, motor, and auditory cortices while largely preserving their existing interactions. Outside the visual system, connectivity changes were more limited and were primarily characterized by selective strengthening of sensorimotor interactions. Together, these findings indicate broad redistribution of cortical connectivity affecting both visual and non-visual systems.

### Projecting (Excitatory) Neurons (Thy1-GCaMP6s)

In Thy1-expressing enucleated mice (**Figure 3A**), cortical connectivity underwent a selective reorganization largely confined to the visual network. V1 preferentially lost connectivity with lateral visual and selected non-visual regions while preserving existing connections and strengthening connectivity with medial visual areas. Likewise, lateral visual areas exhibited selective losses of both intra-visual and cross-modal connectivity. In contrast, medial visual areas (PM, A, and AM) gained connectivity with somatosensory and associative cortices while largely preserving their pre-existing interactions. Outside the visual system, connectivity remained largely preserved, with only limited and spatially restricted changes. Together, these findings indicate that early visual deprivation selectively remodels excitatory cortical networks, primarily through strengthening of medial visual circuits while preserving the overall organization of non-visual systems.

### Inhibitory Neurons (mDLX-GCaMP6s)

In mDLX-expressing enucleated mice (**Figure 3A**), cortical connectivity remained largely preserved following early visual deprivation. V1 exhibited selective losses of connectivity with auditory and trigeminal somatosensory cortices while maintaining its interactions with higher-order visual areas. Likewise, lateral visual cortices displayed only limited alterations, whereas medial visual areas (PM, A, and AM) gained selective connectivity with somatosensory, motor, and associative regions. Outside the visual system, connectivity changes were sparse and largely confined to isolated sensorimotor interactions. Together, these findings indicate that inhibitory cortical networks underwent relatively limited and spatially restricted reorganization in this model of early visual deprivation.

Collectively, these findings demonstrate that the significant interaction between neuronal population and visual status arises from distinct patterns of cortical network reorganization across promoter-defined neuronal populations. Pan-neuronal networks exhibited widespread redistribution, excitatory networks underwent selective remodeling centered on medial visual areas, whereas inhibitory networks remained comparatively preserved.

## Neuronal identity-dependent Reorganization of Cortical Node Influence Induced by Early

### Visual Deprivation

To characterize cell-identity-dependent reorganization of cortical nodal influence following early visual deprivation, we computed nodal efficiency (efficient hubs), connectivity quality (quality hubs), and eigenvector centrality (central hubs) (Lohmann et al. 2010; Rubinov and Sporns 2010). Candidate hubs were identified from the top 15% of each graph metric and evaluated across proportional density thresholds (32–45%). Nodes retained across at least 70%, 80%, or 90% of thresholds were classified as robust, highly robust, or exceptionally robust hubs, respectively (**Figures 4–6**). Hub classifications are presented descriptively to illustrate patterns of nodal influence and were not subjected to independent statistical inference.

**Figure 4.**
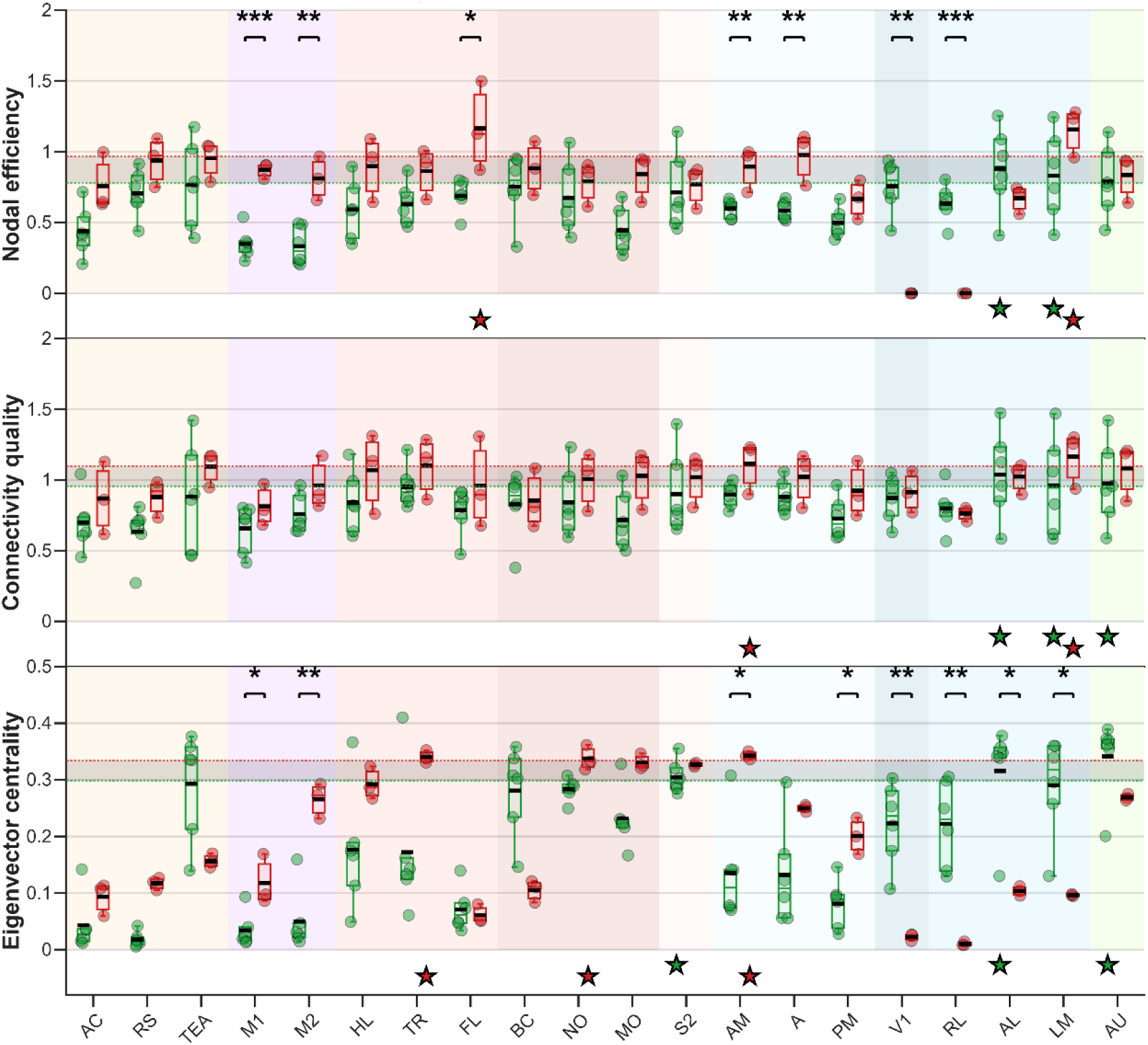
– Effects of Early Visual Deprivation on Cortical Nodal Influence Metrics in Pan-Neuronal Population (hSyn-GCaMP6s) of C57BL/6J Mice. Regional distributions of nodal efficiency, connectivity quality, and eigenvector centrality are shown for sighted (green) and enucleated (red) mice across cortical regions. Metrics were averaged across network density thresholds ranging from 32% to 45%. Background colors indicate the functional classification of each cortical area according to the grouping scheme defined in Figure 1C. Boxplots represent the interquartile range, with whiskers corresponding to the default boxplot distribution range; individual dots represent single animals, and black horizontal bars denote group means. Dashed horizontal lines indicate the top 15% threshold values used for hub identification in each group, and the shaded grey region represents the overlap between sighted and enucleated hub thresholds. Star markers below the x-axis identify robust hub regions for each condition. Significant group differences following FDR-corrected comparisons are indicated by asterisks (* p < 0.05, ** p < 0.01, *** p < 0.001).

**Figure 5.**
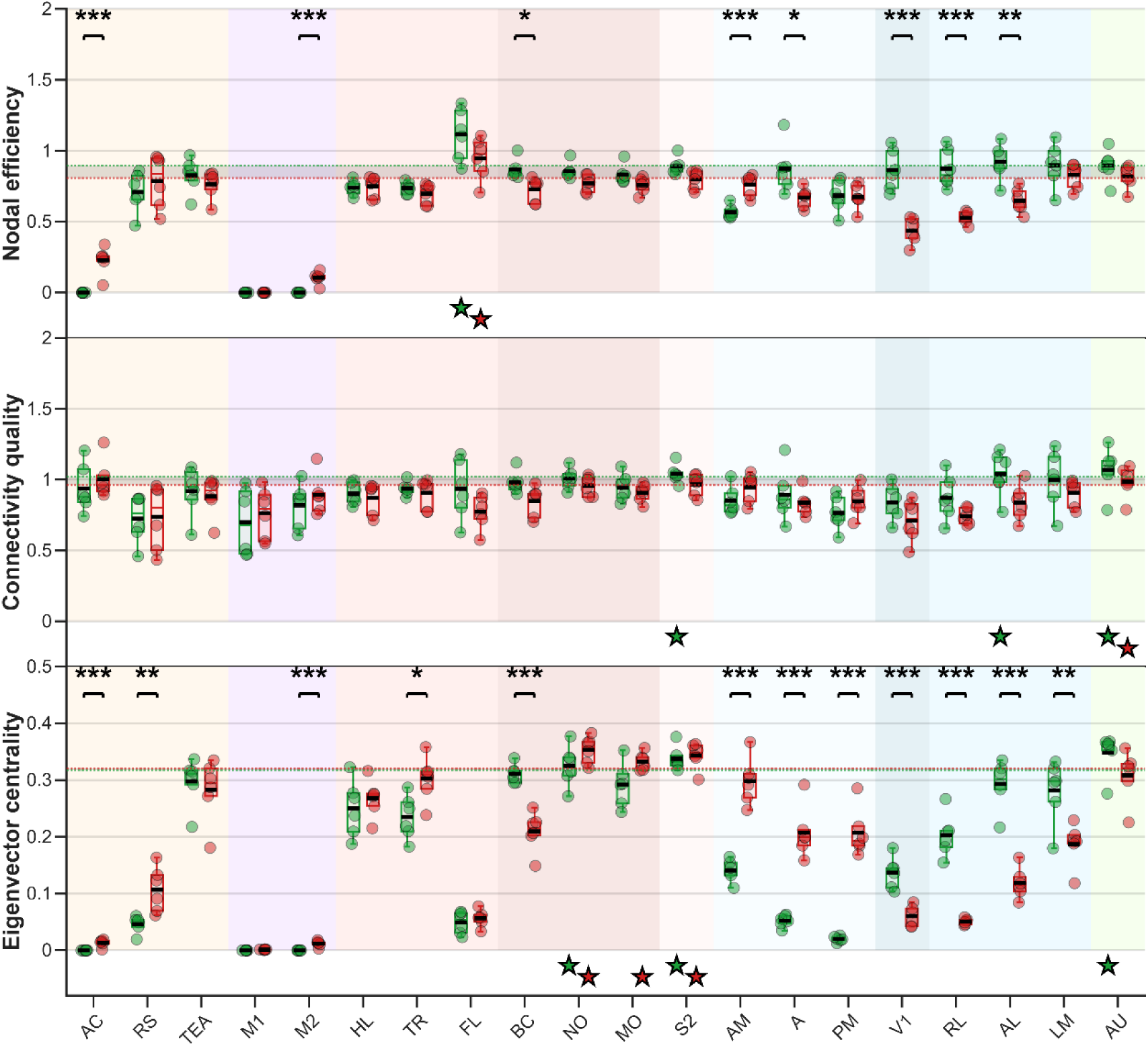
– Effects of Early Visual Deprivation on Cortical Nodal Influence Metrics in Projecting (Excitatory) Neuronal Population (Thy1-GCaMP6s) of C57BL/6J Mice. Regional distributions of nodal efficiency, connectivity quality, and eigenvector centrality are shown for sighted (green) and enucleated (red) mice in the excitatory projecting neuronal population expressing *Thy1-GCaMP6s*. Functional region grouping, hub identification criteria, graphical conventions, and statistical procedures are identical to those described in Figure 4.

**Figure 6.**
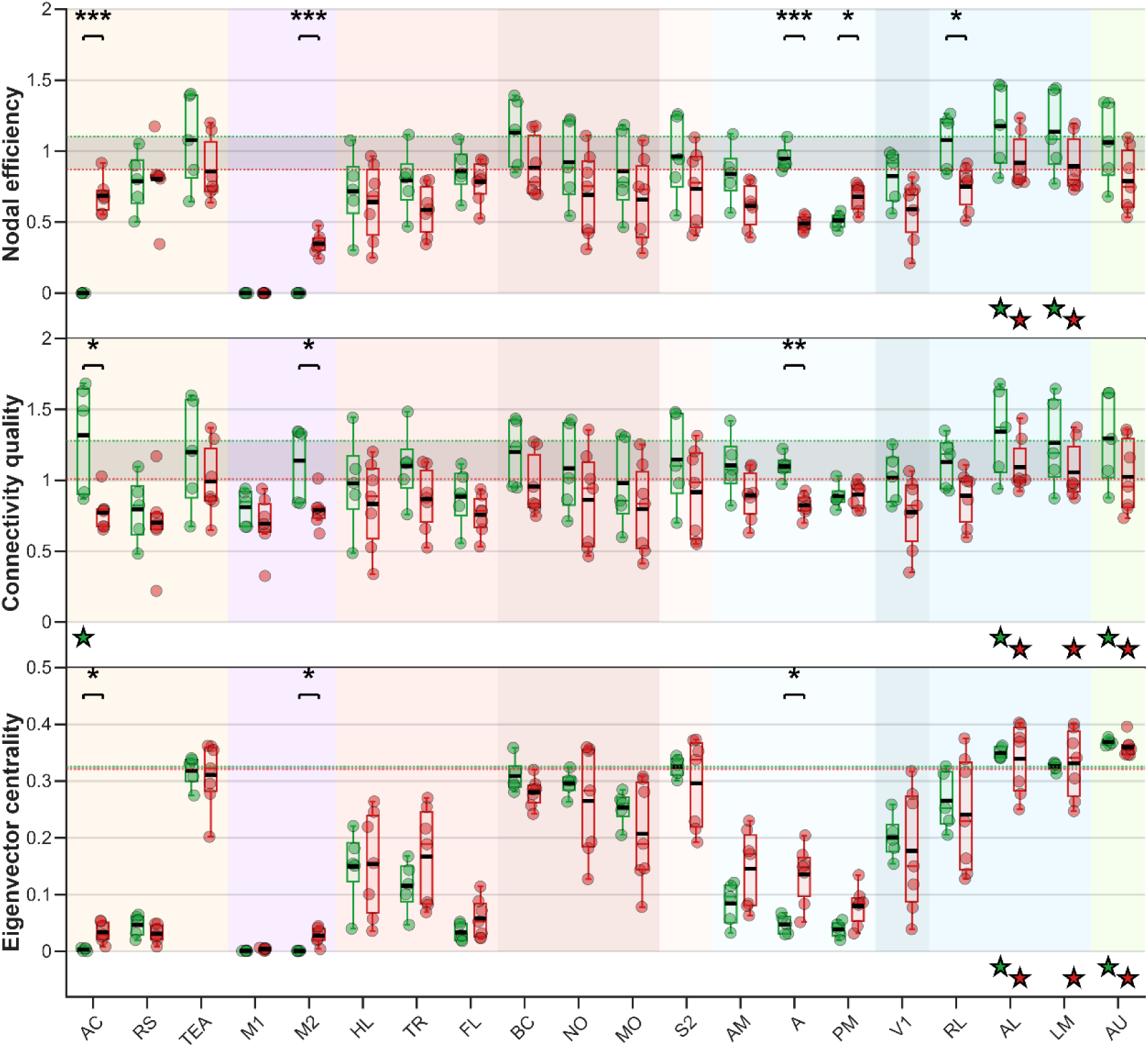
– Effects of Early Visual Deprivation on Cortical Nodal Influence Metrics in Inhibitory Neuronal Population (mDLX-GCaMP6s) of C57BL/6J Mice. Regional distributions of nodal efficiency, connectivity quality, and eigenvector centrality are shown for sighted (green) and enucleated (red) mice in the inhibitory neuronal population expressing *mDLX-GCaMP6s*. Functional region grouping, hub identification criteria, graphical conventions, and statistical procedures are identical to those described in Figure 4.

### Neuronal Identity and Sensory Deprivation Differentially Shape Spontaneous Cortical Network Influence

Multivariate analysis of nodal influence metrics revealed robust effects of visual deprivation across cortical systems, together with a strong modulation by neuronal populations. Significant main effects of visual status were observed across all functional domains, indicating a widespread impact of early enucleation on cortical network organization.

Critically, these effects were not uniform across neuronal populations. To identify the functional systems contributing to these differences while limiting dimensionality, nodal influence metrics were first examined using domain-level multivariate analyses. Significant Promoter × Visual Status interactions were detected across visual, motor, associative, auditory, and lemniscal somatosensory systems, whereas trigeminal somatosensory regions showed only a non-significant trend after permutation correction. Targeted ROI-level multivariate analyses confirmed that these interaction effects were distributed across visual, motor, associative, and somatosensory cortices. Across regions, nodal efficiency and eigenvector centrality accounted for most interaction effects, whereas connectivity quality exhibited more selective population-dependent modulation.

Within the visual system, interaction effects revealed a differential reorganization of the medial and lateral visual streams. Primary visual cortex (V1) exhibited a significant interaction effect on nodal efficiency (F(2,27) = 5.82, p = 0.019, η²p = 0.30). The medial visual stream exhibited the strongest population-dependent modulation, including area A (nodal efficiency: F(2,27) = 38.04, p < 0.001, η²p = 0.74; quality: F(2,27) = 6.94, p = 0.021, η²p = 0.34), AM (nodal efficiency: F(2,27) = 13.05, p < 0.001, η²p = 0.49), and PM (eigenvector centrality: F(2,27) = 15.32, p < 0.001, η²p = 0.53). The lateral visual stream also showed significant interaction effects in eigenvector centrality (AL: F(2,27) = 10.49, p = 0.001, η²p = 0.44; LM: F(2,27) = 7.69, p = 0.006, η²p = 0.36; RL: F(2,27) = 5.72, p = 0.018, η²p = 0.30), indicating that neuronal identity-dependent remodeling involved both visual pathways while preferentially affecting higher-order medial visual regions.

Population-dependent reorganization extended to sensorimotor and associative networks. Lemniscal somatosensory regions exhibited significant modulation of nodal efficiency (FL: F(2,27) = 9.58, p = 0.001, η²p = 0.42), demonstrating that these effects extended beyond visual cortices. Motor regions exhibited the strongest interaction effects observed in the study. M1 showed the largest effect size (nodal efficiency: F(2,27) = 90.43, p < 0.001, η²p = 0.87), together with a significant effect on eigenvector centrality (F(2,27) = 15.33, p < 0.001, η²p = 0.53). M2 similarly exhibited significant modulation across all nodal influence metrics (eigenvector centrality: F(2,27) = 42.89, p = 0.013, η²p = 0.76; nodal efficiency: F(2,27) = 13.88, p < 0.001, η²p = 0.51; quality: F(2,27) = 7.70, p = 0.019, η²p = 0.36), highlighting substantial reorganization of motor network influence. Associative cortices also contributed to these interaction effects, with AC exhibiting significant modulation of nodal efficiency (F(2,27) = 12.22, p < 0.001, η²p = 0.48), indicating that neuronal identity-dependent remodeling extends to higher-order integrative networks.

These findings demonstrate that early visual deprivation differentially remodeled cortical nodal influence according to neuronal population. Consistent with the connectivity and graph-theoretical analyses, this reorganization is characterized by preferential remodeling of the medial relative to the lateral visual stream, together with substantial changes in motor network influence, providing the foundation for the population-specific hub analyses presented below.

### Pan-Neuronal Population (hSyn-GCaMP6s)

Multivariate analysis of nodal influence metrics identified the visual, lemniscal somatosensory, and motor systems as the principal contributors to network reorganization following early visual deprivation within the pan-neuronal (hSyn) population (**Figure 4**).

Within the visual system, nodal influence was redistributed from the lateral toward the medial visual stream. Primary visual cortex (V1) exhibited marked reductions in nodal efficiency (F(1,7) = 45.67, p = 0.001, η²p = 0.87; Δ = –0.76) and eigenvector centrality (F(1,7) = 21.34, p = 0.01, η²p = 0.75; Δ = –0.20). Similar reductions were observed throughout the lateral visual stream, particularly in RL (nodal efficiency: F(1,7) = 69.08, p < 0.001, η²p = 0.91; Δ = –0.63; eigenvector centrality: F(1,7) = 21.54, p = 0.01, η²p = 0.75; Δ = –0.21), with additional decreases in eigenvector centrality in AL (F(1,7) = 14.79, p = 0.01; Δ = –0.21) and LM (F(1,7) = 13.60, p = 0.01; Δ = –0.19). In contrast, the medial visual stream showed increased nodal influence, including area A (nodal efficiency: F(1,7) = 22.54, p = 0.005, η²p = 0.76; Δ = +0.39), AM (nodal efficiency: F(1,7) = 17.43, p = 0.008, η²p = 0.71; Δ = +0.29; eigenvector centrality: F(1,7) = 14.61, p = 0.013, η²p = 0.68; Δ = +0.21), and PM (eigenvector centrality: F(1,7) = 17.70, p = 0.012, η²p = 0.72; Δ = +0.12). Consistent with this redistribution, AL lost its exceptionally robust efficient, quality, and central hub profiles following enucleation, whereas AM emerged as a highly robust quality and exceptionally robust central hub. LM retained highly robust efficient and quality hub classifications across visual conditions.

Beyond the visual system, nodal influence shifted toward somatosensory and motor networks. Within the lemniscal somatosensory system, FL showed increased nodal efficiency (F(1,7) = 12.23, p = 0.017, η²p = 0.64; Δ = +0.48), emerging as an exceptionally stable efficient hub following enucleation. TR and NO also acquired highly robust central hub profiles, whereas S2 and the auditory cortex (AU) no longer retained robust influence hub classifications.

Motor regions exhibited robust increases in network influence. Nodal efficiency increased significantly in both M1 (F(1,7) = 62.74, p < 0.001, η²p = 0.90; Δ = +0.52) and M2 (F(1,7) = 23.36, p = 0.005, η²p = 0.77; Δ = +0.48), accompanied by increased eigenvector centrality (M1: F(1,7) = 11.22, p = 0.018, η²p = 0.62; Δ = +0.08; M2: F(1,7) = 38.39, p = 0.005, η²p = 0.85; Δ = +0.22). Despite these increases, neither motor region consistently met the criteria for robust hub classification across density thresholds.

Together, these findings indicate widespread redistribution of cortical nodal influence, characterized by reduced influence of lateral visual regions and increased influence of medial visual and sensorimotor networks.

### Projecting (Excitatory) Neurons (Thy1-GCaMP6s)

Multivariate analysis of nodal influence metrics identified the primary visual, lateral and medial visual, lemniscal somatosensory, auditory, motor, and associative systems as the principal contributors to network reorganization following early visual deprivation within the excitatory (Thy1) population (**Figure 5**).

Within the visual system, nodal influence underwent a marked redistribution between the lateral and medial visual streams. Primary visual cortex (V1) exhibited marked reductions in nodal efficiency (F(1,10) = 38.58, p < 0.001, η²p = 0.79; Δ = –0.43) and eigenvector centrality (F(1,10) = 33.06, p < 0.001, η²p = 0.77; Δ = –0.08). Similar reductions characterized the lateral visual stream, particularly RL (nodal efficiency: F(1,10) = 36.12, p < 0.001, η²p = 0.78; Δ = –0.35; eigenvector centrality: F(1,10) = 97.13, p < 0.001, η²p = 0.91; Δ = –0.15) and AL (nodal efficiency: F(1,10) = 19.62, p = 0.003, η²p = 0.66; Δ = –0.27; eigenvector centrality: F(1,10) = 73.89, p < 0.001, η²p = 0.88; Δ = –0.17), with additional reductions in LM eigenvector centrality (F(1,10) = 12.14, p = 0.009, η²p = 0.55; Δ = –0.09). In contrast, the medial visual stream displayed increased nodal influence. AM exhibited increased nodal efficiency (F(1,10) = 29.32, p < 0.001, η²p = 0.75; Δ = +0.19) and eigenvector centrality (F(1,10) = 72.71, p < 0.001, η²p = 0.88; Δ = +0.16), whereas PM showed a marked increase in eigenvector centrality (F(1,10) = 119.92, p < 0.001, η²p = 0.92; Δ = +0.19). In contrast, area A exhibited reduced nodal efficiency (F(1,10) = 7.60, p = 0.04, η²p = 0.43; Δ = –0.20) together with increased eigenvector centrality (F(1,10) = 67.48, p < 0.001, η²p = 0.87; Δ = +0.16), revealing dissociation between local efficiency and global network influence. Consistent with these changes, AL functioned as a highly robust quality hub in sighted animals, but no visual region retained robust influence hub classification following enucleation. Conversely, AM acquired a robust provincial and bridging hub profile at the modular level, suggesting increased modular embedding rather than global influence centrality.

Outside visual cortices, neuronal identity-dependent reorganization extended to sensory, motor, and associative networks. Within the somatosensory system, trigeminal regions showed reduced nodal influence (BC nodal efficiency: F(1,10) = 9.98, p = 0.023, η²p = 0.50; Δ = –0.14; eigenvector centrality: F(1,10) = 42.03, p < 0.001, η²p = 0.81; Δ = –0.10), whereas lemniscal regions showed increased integration (TR eigenvector centrality: F(1,10) = 9.15, p = 0.017, η²p = 0.48; Δ = +0.07). FL retained an exceptionally robust efficient hub profile, while MO emerged as an exceptionally robust central hub following enucleation. NO and S2 retained exceptionally robust central hub classifications, although S2 no longer fulfilled robust quality hub criteria after visual deprivation.

ROI-level MANOVA identified significant multivariate reorganization within the auditory cortex (Pillai’s trace = 0.71, F(3,8) = 6.47, p = 0.02, η²p = 0.71), despite the absence of significant univariate effects, indicating distributed multimetric remodeling. AU retained a highly robust quality hub profile across visual conditions but no longer exhibited exceptionally robust central hub classification following enucleation.

Motor and associative cortices also exhibited increased nodal influence. M2 showed significant increases in nodal efficiency (F(1,10) = 36.11, p < 0.001, η²p = 0.78; Δ = +0.10) and eigenvector centrality (F(1,10) = 32.33, p < 0.001, η²p = 0.76; Δ = +0.01), although no motor region fulfilled robust influence hub criteria. Associative regions likewise contributed to this reorganization, with AC exhibiting increased nodal efficiency (F(1,10) = 34.18, p < 0.001, η²p = 0.77; Δ = +0.23) and eigenvector centrality (F(1,10) = 27.21, p < 0.001, η²p = 0.73; Δ = +0.01), while RS showed increased eigenvector centrality (F(1,10) = 12.42, p = 0.009, η²p = 0.55; Δ = +0.06).

Together, these findings indicate selective remodeling of nodal influence centered on medial visual and associative networks.

### Inhibitory Neurons (mDLX-GCaMP6s)

Multivariate analysis of nodal influence metrics identified the lateral and medial visual, lemniscal somatosensory, auditory, motor, and associative systems as the principal contributors to network reorganization within the inhibitory (mDLX) population following early visual deprivation (**Figure 6**). In contrast to the pan-neuronal and excitatory populations, primary visual and trigeminal somatosensory domains did not show significant multivariate modulation.

Within the visual system, nodal influence exhibited comparatively limited redistribution while robust hub organization was preserved. The lateral visual stream showed only selective reductions, confined primarily to RL (nodal efficiency: F(1,10) = 10.35, p = 0.017, η²p = 0.51; Δ = –0.33). Medial visual regions displayed more differentiated responses. Area A exhibited reduced nodal efficiency (F(1,10) = 121.87, p < 0.001, η²p = 0.92; Δ = –0.46) and connectivity quality (F(1,10) = 33.87, p = 0.002, η²p = 0.77; Δ = –0.27), together with increased eigenvector centrality (F(1,10) = 13.18, p = 0.014, η²p = 0.57; Δ = +0.09), indicating reduced local influence but enhanced global integration. PM showed increased nodal efficiency (F(1,10) = 12.11, p = 0.013, η²p = 0.55; Δ = +0.17), whereas AM exhibited significant multivariate modulation (Pillai’s trace = 0.80, F(3,8) = 10.60, p = 0.009, η²p = 0.80) without significant changes in individual nodal metrics. Despite these regional alterations, AL retained exceptionally robust efficient, quality, and central hub classifications, whereas LM retained an exceptionally robust efficient hub profile and acquired exceptionally robust quality and central hub classifications following enucleation.

Outside the visual system, reorganization primarily involved associative and motor networks while preserving stable hub architecture in sensory systems. Within the lemniscal somatosensory system, ROI-level MANOVA identified significant modulation in TR (Pillai’s trace = 0.65, F(3,8) = 4.99, p = 0.048, η²p = 0.65), although no individual nodal metric reached significance, indicating distributed multimetric reorganization. A similar pattern was observed in the auditory cortex (Pillai’s trace = 0.75, F(3,8) = 7.85, p = 0.016, η²p = 0.75), where AU retained highly robust quality and exceptionally stable central hub classifications across visual conditions.

Motor and associative cortices exhibited the most prominent changes outside the visual system. M2 showed increased nodal efficiency (F(1,10) = 105.82, p < 0.001, η²p = 0.91; Δ = +0.35) and eigenvector centrality (F(1,10) = 18.15, p = 0.014, η²p = 0.64; Δ = +0.03), accompanied by reduced connectivity quality (F(1,10) = 9.34, p = 0.036, η²p = 0.48; Δ = –0.35). Similarly, AC exhibited increased nodal efficiency (F(1,10) = 152.68, p < 0.001, η²p = 0.94; Δ = +0.68) and eigenvector centrality (F(1,10) = 14.48, p = 0.014, η²p = 0.59; Δ = +0.03), together with reduced connectivity quality (F(1,10) = 11.94, p = 0.028, η²p = 0.54; Δ = –0.55). Although RS exhibited significant multivariate modulation (Pillai’s trace = 0.82, F(3,8) = 12.26, p = 0.008, η²p = 0.82), no individual nodal metric reached significance. Consistent with these findings, AC no longer retained robust quality hub classification following enucleation, and no associative region acquired robust influence hub status.

Together, these findings indicate that the hub organization of the inhibitory networks was largely preserved despite localized changes in nodal influence.

Collectively, these findings indicate that early visual deprivation induced population-dependent patterns of cortical nodal reorganization. Within pan-neuronal networks, nodal influence was redistributed from lateral toward medial visual circuits and sensorimotor regions. Within excitatory projection networks, remodeling was centered on higher-order visual and associative cortices. Within inhibitory networks, pre-existing hub organization remained largely preserved despite localized changes in nodal influence metrics.

## Spontaneous Cortical Activity Reveals Modular Reorganization Across Pan-Neuronal, Excitatory, and Inhibitory Networks in Early Blind Mice

To characterize modular reorganization following early visual deprivation, Louvain community detection was used to identify functional modules (Blondel *et al*. 2008). Participation coefficient (PC), within-module centrality (WMC), and betweenness centrality (BeC) were computed to identify connector, provincial, and bridging hubs, respectively (Guimera and Nunes Amaral 2005; van den Heuvel and Sporns 2011). Candidate hubs were identified from the top 15% of each graph metric and evaluated across proportional density thresholds (32–45%). Nodes retained across at least 70% of thresholds were considered robust hubs and retained for graphical representation (**Figures 7–9**). Hub classifications were interpreted descriptively and were not subjected to independent statistical inference.

**Figure 7.**
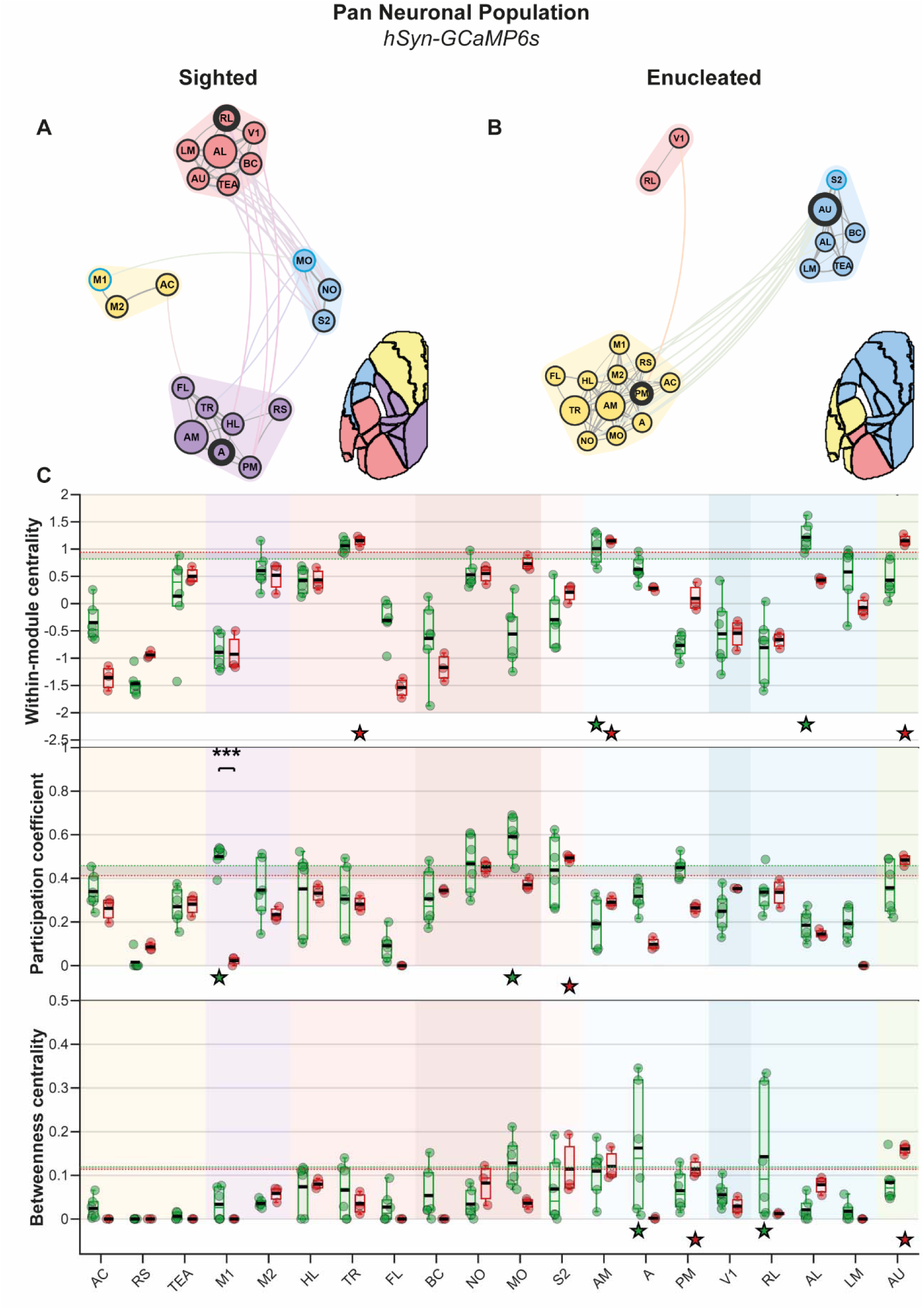
– Network Community Structure and Structural Nodal Metrics in Pan-Neuronal Population (hSyn-GCaMP6s) C57BL/6J Mice. **A.** Community organization and structural hub architecture in sighted mice. Community maps were generated using Louvain modularity consensus analysis across density thresholds (32–45%). Enlarged nodes identify provincial hubs, cyan outline connector hubs, and thick borders bridging hubs. **B.** Community organization and structural hub architecture in enucleated mice displayed using the same graphical conventions and hub-classification criteria as in **panel A**. **C.** Regional distributions of within-module centrality, participation coefficient, and betweenness centrality in sighted (green) and enucleated (red) mice. Background colors indicate functional cortical groupings as defined in Figure 1C. Dashed lines indicate hub-classification thresholds. Star markers below the x-axis identify robust provincial, connector, or bridging hubs. Significant between-group differences following FDR-corrected comparisons are indicated by asterisks (* p < 0.05, ** p < 0.01, *** p < 0.001).

**Figure 8.**
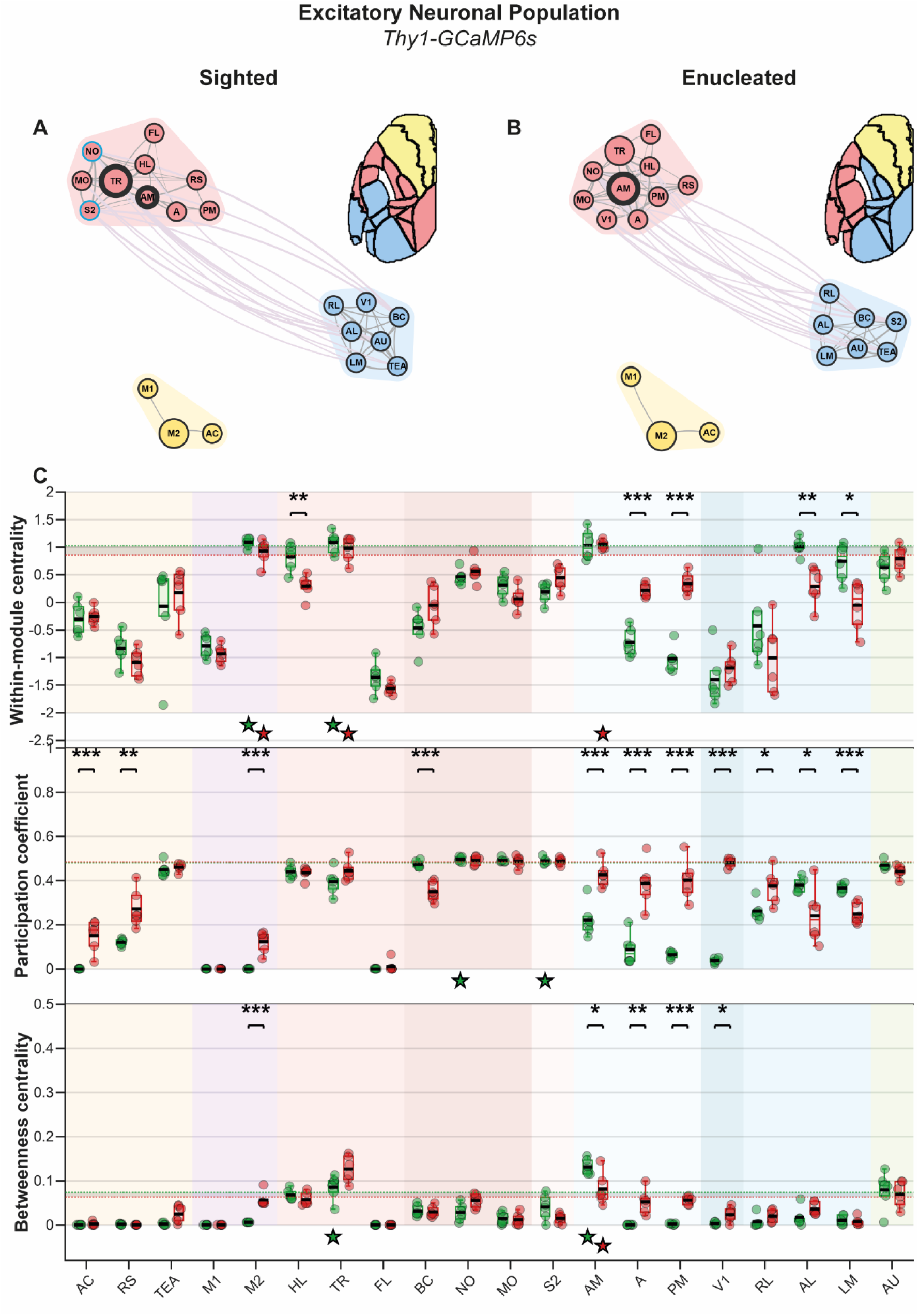
– Network Community Structure and Structural Nodal Metrics in Projecting (Excitatory) Neuronal Population (Thy1-GCaMP6s) C57BL/6J Mice. **A.** Community organization and structural hub architecture in sighted mice. **B.** Community organization and structural hub architecture in enucleated mice using the same graphical conventions and hub-classification criteria as in **panel A**. **C.** Regional distributions of within-module centrality, participation coefficient, and betweenness centrality in sighted (green) and enucleated (red) mice. Graphical conventions are identical to Figure 7.

**Figure 9.**
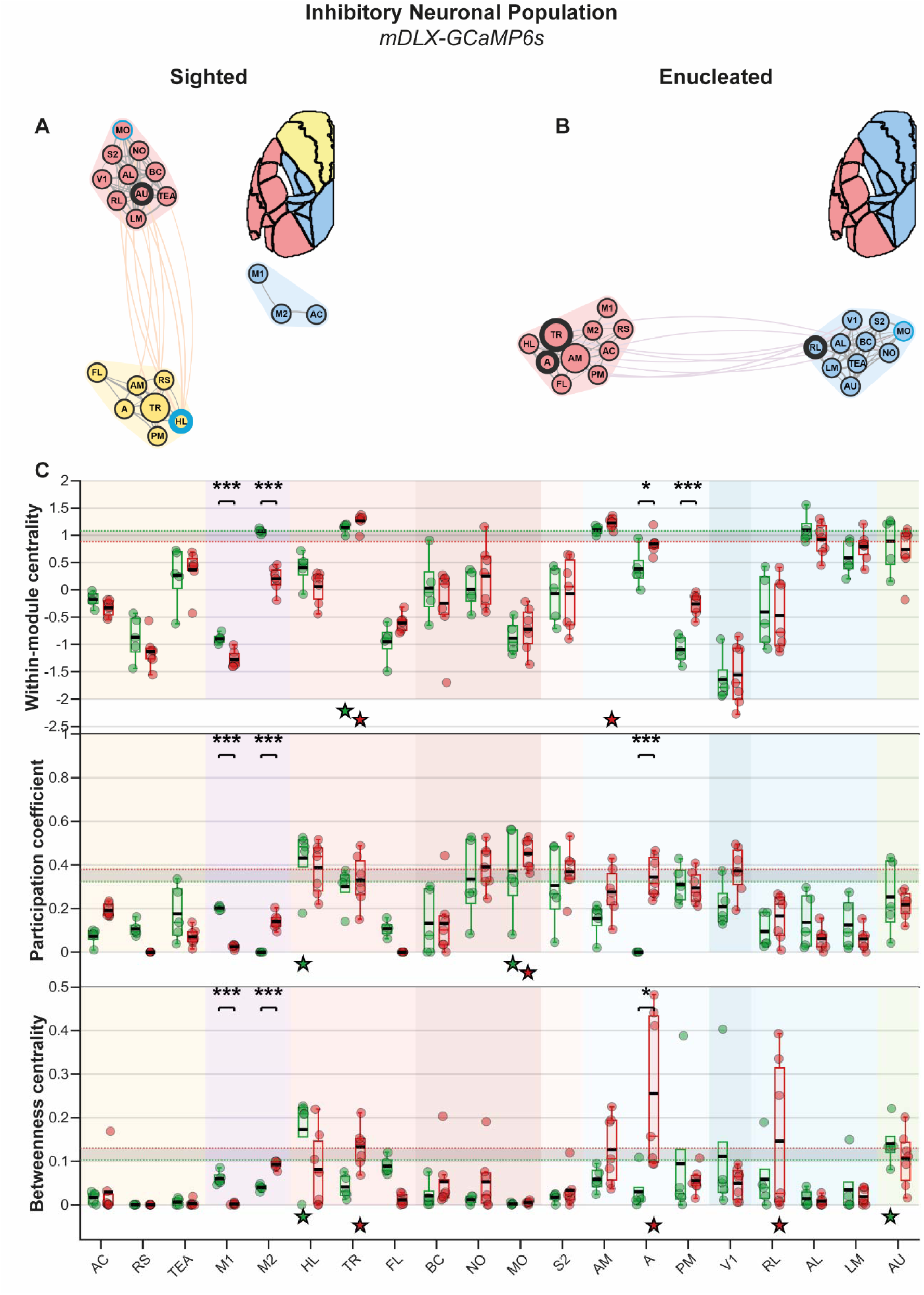
– Network Community Structure and Structural Nodal Metrics in Inhibitory Neuronal Population (mDLX-GCaMP6s) C57BL/6J Mice. **A.** Community organization and structural hub architecture in sighted mice. **B.** Community organization and structural hub architecture in enucleated mice using the same graphical conventions and hub-classification criteria as in **panel A**. **C.** Regional distributions of within-module centrality, participation coefficients, and betweenness centrality in sighted (green) and enucleated (red) mice. Graphical conventions are identical to Figure 7.

### Neuronal Identity and Visual Status Shape Module Centrality and Network Integration

Multivariate analysis of nodal structure metrics revealed robust effects of neuronal population and visual deprivation across cortical systems, together with significant neuronal identity-dependent modulation of modular organization. Significant main effects of visual status were observed across primary, lateral, and medial visual domains as well as motor regions, whereas neuronal population additionally influenced visual, somatosensory, auditory, associative, and motor systems.

Significant *Promoter × Visual Status* interactions were detected across primary and medial visual, auditory, motor, and associative cortices. ROI-level analyses showed that PC accounted for most interaction effects, whereas WMC and BeC exhibited more selective regional alteration.

Within the visual system, interaction effects were most prominent in higher-order medial visual regions. Primary visual cortex (V1) exhibited a significant interaction effect on PC (F(2,27) = 15.89, p < 0.001, η²p = 0.54). Medial visual areas showed the strongest population-dependent modulation, including area A (PC: F(2,27) = 39.74, p < 0.001, η²p = 0.74; WMC: F(2,27) = 20.81, p < 0.001, η²p = 0.61; BeC: F(2,27) = 7.70, p = 0.011, η²p = 0.36) and PM (PC: F(2,27) = 43.24, p < 0.001, η²p = 0.76; WMC: F(2,27) = 5.40, p = 0.021, η²p = 0.29), indicating neuronal identity-dependent remodeling of modular integration within higher-order visual networks.

Beyond the visual system, motor regions exhibited the strongest interaction effects. M1 showed significant interactions for PC (F(2,27) = 211.65, p < 0.001, η²p = 0.94) and BeC (F(2,27) = 8.40, p = 0.011, η²p = 0.38), whereas M2 exhibited significant interactions across all three modular metrics (PC: F(2,27) = 9.99, p < 0.001, η²p = 0.42; WMC: F(2,27) = 10.75, p = 0.001, η²p = 0.44; BeC: F(2,27) = 5.34, p = 0.028, η²p = 0.28), identifying motor regions as major contributors to neuronal identity-dependent remodeling of cortical modular architecture. Associative regions also exhibited significant interaction effects, including RS (PC: F(2,27) = 26.86, p < 0.001, η²p = 0.67; WMC: F(2,27) = 5.84, p = 0.020, η²p = 0.30), AC (PC: F(2,27) = 12.76, p < 0.001, η²p = 0.49; WMC: F(2,27) = 13.48, p < 0.001, η²p = 0.50), and TEA (BeC: F(2,27) = 6.64, p = 0.015, η²p = 0.33), indicating that modular reorganization extended beyond sensory systems.

These findings demonstrate that early visual deprivation differentially remodeled cortical modular architecture across promoter-defined neuronal populations, with altered intermodular communication representing the predominant signature of network reorganization.

### Pan-Neuronal Population (hSyn-GCaMP6s)

Multivariate analysis of nodal structure metrics identified the motor domain as the only functional system showing significant modular reorganization following early visual deprivation within the pan-neuronal (hSyn) population (**Figure 7**). Accordingly, ROI-level statistical inference focused on this domain. Patterns observed in the visual, somatosensory, auditory, and associative systems are also described to provide a broader view of potential network reorganization, while changes in hub classification within these domains should be interpreted descriptively because they did not meet the hierarchical criterion for ROI-level inference.

Within the visual system, modular reorganization primarily reflected redistribution of provincial and bridging hub organization between lateral and medial visual regions. AM retained a robust provincial hub profile across visual conditions, whereas PM emerged as a robust bridging hub following enucleation. In contrast, area A and RL no longer retained robust bridging hub classification, and AL lost its robust provincial hub profile.

Outside the motor system, visual conditions were associated with distinct descriptive patterns of hub organization. Although these differences did not reach significance at the domain level, they provide additional context for the significant motor findings. Within the somatosensory system, S2 emerged descriptively as a robust connector hub following enucleation, whereas MO no longer met robust connector hub criteria and TR acquired a robust provincial hub profile. AU also emerged as both a highly robust provincial and bridging hub, a pattern descriptively consistent with greater modular integration of the auditory cortex.

Motor cortex exhibited the only significant quantitative reorganization. M1 showed a marked reduction in PC (F(1,7) = 192.19, p < 0.001, η²p = 0.96; Δ = –0.48), indicating reduced intermodular connectivity. Consistent with this change, M1 no longer retained robust connector hub classification following visual deprivation.

Together, these findings indicate that significant modular reorganization within pan-neuronal networks was concentrated in the motor system, against a broader descriptive background of altered hub-classification patterns across cortical systems.

### Projecting (Excitatory) Neuronal Population (Thy1-GCaMP6s)

Multivariate analysis of nodal structure metrics identified the primary, lateral, and medial visual, associative, motor, and somatosensory systems as the principal contributors to modular reorganization following early visual deprivation within the excitatory (Thy1) population (**Figure 8**). ROI-level analyses further revealed widespread remodeling of modular organization across visual, motor, associative, and somatosensory cortices. Within the visual system, modular organization underwent extensive redistribution between the lateral and medial visual streams. Primary visual cortex (V1) exhibited marked increases in PC (F(1,10) = 2386.80, p < 0.001, η²p = 0.996; Δ = +0.44) and BeC (F(1,10) = 8.28, p = 0.040, η²p = 0.45; Δ = +0.02). The lateral visual stream generally showed reduced modular integration, with LM exhibiting reduced PC (F(1,10) = 33.62, p < 0.001, η²p = 0.77; Δ = –0.12) and WMC (F(1,10) = 12.63, p = 0.013, η²p = 0.56; Δ = –0.79), and AL showing reduced WMC (F(1,10) = 23.30, p = 0.003, η²p = 0.70; Δ = –0.72) and PC (F(1,10) = 6.96, p = 0.027, η²p = 0.41; Δ = – 0.14). In contrast, RL exhibited increased PC (F(1,10) = 10.15, p = 0.012, η²p = 0.50; Δ = +0.11). The medial visual stream displayed robust multimetric increases. PM exhibited increased PC (F(1,10) = 82.66, p < 0.001, η²p = 0.89; Δ = +0.34), WMC (F(1,10) = 126.94, p < 0.001, η²p = 0.93; Δ = +1.36), and BeC (F(1,10) = 204.54, p < 0.001, η²p = 0.95; Δ = +0.05). Area A similarly exhibited increased PC (F(1,10) = 36.60, p < 0.001, η²p = 0.79; Δ = +0.30), WMC (F(1,10) = 71.06, p < 0.001, η²p = 0.88; Δ = +0.94), and BeC (F(1,10) = 21.36, p = 0.004, η²p = 0.68; Δ = +0.05). AM showed increased PC (F(1,10) = 29.29, p < 0.001, η²p = 0.75; Δ = +0.21) together with reduced BeC (F(1,10) = 9.24, p = 0.037, η²p = 0.48; Δ = –0.05). Consistent with these changes, AM retained a highly robust bridging hub profile and additionally emerged as a robust provincial hub following enucleation.

Beyond the visual system, modular remodeling extended to somatosensory, motor, and associative networks. Trigeminal somatosensory cortex (BC) showed reduced PC (F(1,10) = 42.78, p < 0.001, η²p = 0.81; Δ = –0.12), whereas dorsal somatosensory cortex (HL) exhibited reduced WMC (F(1,10) = 17.72, p = 0.005, η²p = 0.64; Δ = –0.53). Consistent with these changes, NO and S2 no longer fulfilled robust connector hub criteria, whereas TR retained a highly robust provincial hub profile while losing robust bridging hub classification. No significant auditory domain-level modulation was detected.

Motor and associative cortices also exhibited enhanced modular integration. M2 showed increased PC (F(1,10) = 42.09, p < 0.001, η²p = 0.81; Δ = +0.12) and BeC (F(1,10) = 53.14, p < 0.001, η²p = 0.84; Δ = +0.05), while maintaining an exceptionally stable provincial hub profile across visual conditions. Associative cortices similarly exhibited increased PC in AC (F(1,10) = 27.67, p < 0.001, η²p = 0.73; Δ = +0.15) and RS (F(1,10) = 18.45, p = 0.002, η²p = 0.65; Δ = +0.15).

Together, these findings indicate that excitatory projection networks underwent extensive remodeling of modular organization, characterized by redistribution from lateral toward medial visual circuits together with coordinated reorganization of sensorimotor and associative networks.

### Inhibitory Neuronal Population (mDLX-GCaMP6s)

Multivariate analysis of nodal structure metrics identified the medial visual and motor systems as the principal contributors to modular reorganization following early visual deprivation within the inhibitory (mDLX) population (**Figure 9**). Accordingly, ROI-level statistical inference focused on these two domains. Patterns observed in the lateral visual, somatosensory, auditory, and associative systems are also described to provide a broader view of potential network reorganization, while changes in hub classification within these domains should be interpreted descriptively because they did not meet the hierarchical criterion for ROI-level inference.

Within the visual system, modular reorganization was largely confined to medial visual regions. Area A exhibited increased PC (F(1,10) = 65.87, p < 0.001, η²p = 0.87; Δ = +0.34), WMC (F(1,10) = 9.06, p = 0.013, η²p = 0.48; Δ = +0.46), and BeC (F(1,10) = 7.36, p = 0.029, η²p = 0.42; Δ = +0.23), indicating enhanced modular integration. PM similarly exhibited increased WMC (F(1,10) = 43.73, p < 0.001, η²p = 0.81; Δ = +0.83). Although lateral visual regions did not exhibit significant domain-level modulation, RL emerged as a robust bridging hub following enucleation. Area A likewise acquired a robust bridging hub profile, whereas AM emerged as a robust provincial hub.

Hub organization outside the significantly modulated medial visual and motor domains appeared largely preserved, although several descriptive changes in regional hub classification suggested more subtle patterns of network reorganization. While these patterns were not assessed using ROI-level statistical inference, they provide additional context for the significant domain-level findings. Within the somatosensory system, MO retained a highly robust connector hub profile across visual conditions, whereas TR retained a highly robust provincial hub profile and additionally emerged as a robust bridging hub following enucleation. Conversely, HL no longer met robust connector or bridging hub criteria. No significant auditory or associative domain-level modulation was detected, although AU descriptively no longer met the criteria for robust bridging hub classification.

Motor cortex exhibited the strongest modular reorganization. M1 showed marked reductions in PC (F(1,10) = 1065.49, p < 0.001, η²p = 0.99; Δ = –0.18), BeC (F(1,10) = 102.67, p < 0.001, η²p = 0.91; Δ = –0.06), and WMC (F(1,10) = 24.94, p < 0.001, η²p = 0.71; Δ = –0.38), indicating reduced modular integration. In contrast, M2 exhibited increased PC (F(1,10) = 74.49, p < 0.001, η²p = 0.88; Δ = +0.14) and BeC (F(1,10) = 129.27, p < 0.001, η²p = 0.93; Δ = +0.05), together with reduced WMC (F(1,10) = 75.43, p < 0.001, η²p = 0.88; Δ = –0.87), indicating selective redistribution of modular roles within the motor network.

Together, these findings indicate that inhibitory networks underwent relatively restricted modular remodeling, primarily involving medial visual and motor systems while preserving much of their pre-existing hub organization.

Descriptive comparison of the significant within-population effects and hub-classification patterns indicated distinct forms of modular reorganization across promoter-defined neuronal populations. Thy1 networks showed significant effects across the largest number of functional domains, whereas significant effects in mDLX networks were restricted to medial visual and motor domains and those in hSyn networks to the motor domain. Descriptive hub-classification patterns further suggested substantial redistribution of modular hub identities in hSyn networks, widespread redistribution of connector and provincial hubs in Thy1 networks, and comparatively preserved hub architecture in mDLX networks. Across populations, participation coefficient accounted for most of the significant regional effects, suggesting that early visual deprivation predominantly altered communication between functional modules rather than uniformly reorganizing within-module organization.

## Discussion

The aim of this study was to investigate how early-onset blindness reshapes the large-scale organization of spontaneous cortical functional networks. By examining resting-state activity in projecting (excitatory), inhibitory, and pan-neuronal populations, we show that early visual deprivation induced both shared and neuronal identity-dependent patterns of network reorganizations. Across populations, cortical connectivity shifted from lateral toward medial visual circuits, whereas the magnitude and topological expression of this reorganization differed markedly between neuronal populations.

Previous investigations have primarily described mesoscopic cortical connectivity in excitatory Thy1 or CaMKIIa expressing populations (Vanni and Murphy 2014; Vanni *et al*. 2017; Xiao et al. 2017; Cramer et al. 2019). By contrast, only one study has examined resting-state functional connectivity of inhibitory neuronal populations (PV, SST, VIP) (O’Connor *et al*. 2022). Here, we extend these observations by comparing projection (Thy1), inhibitory (mDLX), and pan-neuronal (hSyn) networks following early visual deprivation. This population-resolved approach provides new insight into how neuronal identity shapes large-scale cortical plasticity after sensory loss.

## Similarities Between Neuronal Populations

In sighted controls, all neuronal populations exhibited a modular architecture supporting large-scale cortical network organization. Within this general framework, however, excitatory and inhibitory networks displayed distinct configurations of connector, bridging, and provincial hubs, indicating complementary contributions to mesoscale network architecture. Following early blindness, despite their distinct cellular identities and connectivity rules (Karnani et al. 2014; Harris and Shepherd 2015; Cardin 2018; Ma et al. 2021), excitatory and inhibitory networks exhibited convergent mesoscale reorganization patterns, including increased prominence of medial higher visual regions and reduced influence of primary and lateral visual areas, although the extent and topological expression of this reorganization differed substantially between neuronal populations.

In early visually deprived mice, all neuronal populations exhibited convergent network-level adjustments. Consistent with the functional connectivity and nodal influence analyses, all neuronal populations exhibited reduced connectivity and influence within V1 and the lateral higher visual areas (AL, LM, RL), whereas medial higher visual areas (AM, PM, A) and associative cortices (AC, RS) gained functional connectivity and nodal influence following early blindness.

Across neuronal populations, regional alterations in participation coefficient indicated reorganization of intermodular communication following early blindness. In the excitatory network, this increase was widely distributed across cortical territories, whereas in the inhibitory network it remained more spatially confined, predominantly affecting frontal regions such as the motor cortices. These results suggest complementary modes of intermodular communication: excitatory populations support widespread communication between functional modules, whereas inhibitory populations contribute to a more focal, frontal-centered coordination of cortical networks. Together, these findings indicate that the shared reorganization observed across neuronal populations primarily reflects redistribution of communication toward medial associative circuits rather than a generalized increase or decrease in cortical connectivity.

This redistribution was particularly prominent within the posterior parietal cortex (PPC), notably areas A and AM, together with the retrosplenial cortex, which consistently gained functional connectivity, nodal influence, and modular hub organization across neuronal populations. Anatomically positioned at the interface between visual, auditory, and somatosensory systems (Gilissen and Arckens 2021), the PPC is well suited to integrate multisensory information. Its increased influence is therefore consistent with its established roles in spatial navigation, sensorimotor integration, spatial attention, and transformations between egocentric and allocentric reference frames (Whitlock 2017). Importantly, this redistribution was consistently observed across these complementary graph-theoretical analyses, indicating that it represents a robust feature of cortical network reorganization rather than a metric-specific observation.

This redistribution from lateral toward medial higher visual areas is consistent with current models of higher visual cortex development. Higher visual areas undergo progressive functional specialization during postnatal development (Murakami et al. 2017), that this specialization depends on patterned binocular visual experience (Salinas et al. 2021), and that distinct higher visual streams exhibit different developmental trajectories (Smith et al. 2017). Moreover, higher visual areas remain highly susceptible to experience-dependent remodeling following altered visual input (Craddock et al. 2023). Collectively, these studies provide a developmental framework that is consistent with the large-scale network reorganization observed here.

These convergent reconfigurations align with previous reports of widespread remodeling in vision-deprived cortices, including alterations in the excitatory–inhibitory balance (Petrus *et al*. 2015; Rączy *et al*. 2022; Ossandón et al. 2023) and enhanced engagement of associative cortices during cross-modal processing (Collignon et al. 2011; Bedny 2017). Notably, the increased influence of the medial HVAs and associative hubs (RS, PM, PPC) observed here mirrors effective connectivity patterns described in early blind humans during complex tasks involving heteromodal sensory processing (Röder *et al*. 2002; Fiehler et al. 2009). This cross-species convergence supports the view that early sensory experience contributes to cortical network organization, leading to sustained integration and functional communication even in the absence of a primary sensory input.

Taken together, these nodal and hub-level reorganizations indicate that cortical resilience emerges from the redistribution of influential roles toward associative and parietal hubs. This interpretation is conceptually consistent with models of connectome robustness (Rubinov and Sporns 2010; van den Heuvel and Sporns 2011), where reallocation of hub influence supports the preservation of global integration despite structural perturbation.

## Stream-Specific Remodeling

Across all neuronal populations, early visual deprivation induced a pronounced stream-specific reorganization of cortical networks. Functional connectivity, nodal influence, and modular organization consistently declined within the lateral higher visual areas, whereas medial higher visual areas preserved or strengthened their integrative roles. This dissociation mirrors human findings showing that, under congenital blindness, ventral occipitotemporal regions remain functionally involved in object and category recognition, while dorsal occipitoparietal regions maintain spatial and sensorimotor processing functions (Bonino et al. 2008; Striem-Amit et al. 2012). Structurally, this dissociation between visual streams was also observed, with blind individuals exhibiting cortical thinning and gray-matter reorganization in ventral territories, with relative preservation of dorsal areas (Qin et al. 2013; Reislev et al. 2016).

This consistent redistribution across multiple graph-theoretical analyses suggests that the lateral and medial visual streams differ fundamentally in their capacity to reorganize following early visual deprivation. Mechanistically, this asymmetry may reflect the distinct thalamocortical and corticothalamic architecture of the murine lateral posterior (LP) nucleus, the homolog of the primate pulvinar (Kaas and Lyon 2007; Bennett et al. 2019; Juavinett et al. 2020). The LP is subdivided into regions with complementary inputs and outputs; its posterior subdivisions receive dominant driving input from the superior colliculus and provide strong feed-forward projections to lateral HVAs (Zhou et al. 2017; Bennett *et al*. 2019; Leow et al. 2022). Consequently, the loss of visual drive through this tecto–thalamo–cortical route could explain the reduced connectivity, nodal influence, and modular integration observed in lateral higher visual areas. In contrast, anterior and medial LP subdivisions receive substantial corticothalamic input from associative and somato-motor cortices (Roth et al. 2016; Bennett *et al*. 2019; Leow *et al*. 2022), and project to medial HVAs, embedding them within cortico–LP–cortical loops that integrate multimodal and feedback information.

Within this framework, the preserved or enhanced role of medial higher visual areas across functional connectivity, nodal influence, and modular analyses is consistent with potentiation of multisensory cortico–LP–cortical circuits. Experimental studies support this interpretation. Lateral HVAs rely predominantly on superior colliculus–LP input (Beltramo and Scanziani 2019), whereas medial HVAs receive proportionally greater associative corticothalamic input (Roth *et al*. 2016; Leow *et al*. 2022). Following early visual deprivation, this circuitry may preferentially propagate multisensory and top-down signals through cortico–thalamo–cortical loops, thereby preserving the integrative role of medial higher visual areas despite the loss of retinal input. Although retinal input is eliminated in the present model, extrageniculo-cortical pathways (Tohmi et al. 2014; Zhou *et al*. 2017) may provide an anatomical substrate supporting this large-scale reorganization.

## Differential Effect of Early Blindness on Specific Neuronal Populations

While convergent reorganization patterns were observed across neuronal populations, early blindness induced neuronal identity-dependent network alterations that reflected the distinct functional roles of excitatory and inhibitory systems.

In excitatory networks, reorganization was predominantly centered on the visual system. Across functional connectivity, nodal influence, and modular analyses, alterations were largely confined to visual cortices, particularly the redistribution from lateral toward medial higher visual areas. This spatial specificity indicates that excitatory networks preferentially reorganize within unimodal visual domains, highlighting their sensitivity to the loss of afferent visual activity. This focal pattern is in line with previous mesoscale imaging studies showing that excitatory networks are organized around topographically restricted functional modules (Vanni and Murphy 2014; Vanni *et al*. 2017; Xiao *et al*. 2017).

In contrast, inhibitory networks exhibited a distinct spatial pattern of functional connectivity reorganization. Unlike pan-neuronal and excitatory networks, whose alterations predominantly involved visual cortices, connectivity changes in inhibitory networks were distributed across multiple functional systems, including auditory, somatosensory, motor, and associative regions. This broader anatomical distribution suggests that inhibitory circuits contribute to coordinating large-scale communication across cortical systems, consistent with their established role in regulating both local synchrony and long-range cortical interactions (Sohal et al. 2009; Isaacson and Scanziani 2011). Consistent with this interpretation, these distributed connectivity changes contrasted with the comparatively modest alterations observed in nodal influence and modular organization, suggesting that inhibitory circuits primarily contribute to preserving the overall topology of cortical networks while selectively adjusting interregional communication during sensory deprivation (Karnani *et al*. 2014; Petrus *et al*. 2015).

Finally, the pan-neuronal network, which integrates both excitatory and inhibitory activity, did not exhibit a simple additive pattern of connectivity changes. Instead, it revealed a nonlinear and synergistic reorganization, suggesting that excitation and inhibition interact dynamically to sustain cortical communication (Okun and Lampl 2008; Karnani et al. 2016; Sadeh and Clopath 2021). This interpretation is consistent with the weaker global topological effects observed in the pan-neuronal analyses despite widespread redistribution of individual connections.

This apparent synergy, however, may partly reflect an important methodological consideration of the mDLX promoter, which labels a mixed population of PV+, SST+, and VIP+ GABAergic interneurons. These GABAergic subtypes play distinct but complementary roles in regulating network excitability and stability (Karnani *et al*. 2014; Karnani *et al*. 2016; Tremblay et al. 2016). As such, the inhibitory contribution observed in the pan-neuronal network likely represents the aggregate influence of multiple inhibitory mechanisms, yielding a composite yet functionally coherent form of reorganization rather than a linear sum of excitatory and inhibitory effects.

Together, these findings demonstrate that early visual deprivation remodeled cortical networks through neuronal identity-dependent patterns. Within excitatory projection networks, deprivation-related topological reorganization was largely centered on visual circuits and involved a redistribution from lateral toward medial higher visual areas. Within inhibitory networks, global topology remained largely preserved despite spatially distributed connectivity changes, consistent with selective adjustment of interregional communication. Considered together, these promoter-specific results support the interpretation that large-scale cortical plasticity emerges from dynamic interactions between excitatory and inhibitory network activity rather than from the independent reorganization of either neuronal population alone.

## Projecting (Excitatory; Thy1) Neuronal Population: Vision-Centric Functional Connectivity Vulnerability

Early visual deprivation induced pronounced reorganization within the excitatory network, characterized by selective alterations in global topology together with widespread redistribution of nodal influence and modular organization. These findings indicate substantial remodeling of long-range cortical organization among glutamatergic cortical neurons, despite relatively localized changes in functional connectivity. This pattern is consistent with the loss of early excitatory visual drive, which deprives the cortical hierarchy of its principal sensory input, as well as with the known sensitivity of global network topology to disruption of excitatory hub regions (Tu et al. 2021). Indeed, experimental inactivation of cortical hubs in awake rodents produces network-wide reductions in global efficiency and modularity, accompanied by the emergence of compensatory integrative hubs such as the retrosplenial cortex (Tu *et al*. 2021).

Consistent with this topological reorganization, in the absence of early visual input, our data reveal redistribution of network integration accompanied by shifts in nodal influence: primary and lateral visual areas lost their dominant role, whereas medial HVAs and RS increased their centrality. The prominent emergence of medial higher visual regions, particularly A, AM, and PM, supports the interpretation that posterior parietal-associated circuits become alternative integrative hubs following early visual deprivation. These medial HVAs and RS, which have extensive reciprocal projections to V1 in sighted mice (Stehberg et al. 2014; Morimoto et al. 2021), exhibited increased influence, suggesting that in early blindness, cortical integration may increasingly rely on these preexisting feedback and associative pathways rather than on direct thalamocortical drive (Lee 2022).

At the modular level, V1 exhibited increased alignment with medial higher visual areas and the retrosplenial cortex, indicating reorganization of its role within the cortical modular architecture. Consistent with the functional connectivity analyses, this modular reorganization was accompanied by enhanced coupling between visual and somatosensory-associative regions, suggesting that the loss of visual input increases the influence of preexisting multisensory pathways and allows cortical communication to be maintained through nonvisual projections (Henschke et al. 2018).

Functionally, this reorganization aligns with models of activity-dependent synaptic plasticity describing strengthening of heteromodal and feedback pathways alongside the weakening of deprived excitatory circuits. Long-range excitatory synapses are known to undergo both short-term potentiation and homeostatic scaling following sensory deprivation (Goel and Lee 2007; Keck et al. 2013; Lee and Whitt 2015). This may provide a mechanistic framework for the observed asymmetric reorganization in which the regions most dependent on direct visual drive lose influence, whereas associative and medial HVAs strengthen their integrative roles through feedback and cortico-cortical connectivity.

These findings are consistent with anatomical and imaging studies reporting large-scale reallocation of excitatory networks following early visual deprivation. In mice, deprivation leads to structural and functional reorganization of intracortical connectivity (Petrus *et al*. 2015), while human neuroimaging studies demonstrate enhanced occipital connectivity with associative and sensorimotor cortices in early blind individuals (Shimony et al. 2006; Shu et al. 2009; Collignon *et al*. 2011; Striem-Amit *et al*. 2012; Bock and Fine 2014; Bauer et al. 2017).

Collectively, these findings demonstrate that early visual deprivation profoundly reorganized excitatory cortical networks. Functional connectivity, nodal influence, and modular organization consistently shifted from primary and lateral visual cortices toward medial higher visual and associative regions, particularly medial higher visual areas and the retrosplenial cortex. This redistribution is consistent with weakening of deprived feedforward visual circuits together with strengthening of preexisting heteromodal and feedback pathways, allowing visual regions to remain integrated within large-scale cortical networks despite the absence of retinal input. These observations suggest that excitatory cortical circuits are particularly sensitive components of the network reorganization associated with early sensory deprivation.

## Inhibitory (mDLX) Neuronal Population: Stable Network Topology Despite Distributed Connectivity Reorganization

At the global level, inhibitory networks exhibited limited topological reorganization following early visual deprivation. None of the global network metrics (global efficiency, modularity, or connectivity strength) changed significantly, indicating relative preservation of large-scale cortical organization despite redistributed functional connectivity. Unlike excitatory networks, whose reorganization remained largely centered on visual cortices, connectivity changes in inhibitory networks were distributed across multiple functional systems, including auditory, somatosensory, motor, and associative regions. This combination of distributed connectivity changes with relatively preserved global topology is consistent with the established role of inhibitory neurons in regulating local circuit dynamics and coordinating interregional communication rather than driving large-scale cortical reorganization (Oswald et al. 2009; Avermann et al. 2012). Because inhibitory activity is embedded within the long-range architecture established by excitatory projections (Harris and Shepherd 2015), preservation of inhibitory network topology despite redistributed connectivity may reflect maintenance of the underlying excitatory scaffold while selectively adjusting communication between cortical areas. This interpretation is consistent with the established role of inhibitory circuits in regulating cortical gain and maintaining excitation–inhibition balance, thereby stabilizing large-scale network dynamics despite changes in interregional communication (Haider and McCormick 2009; Sohal *et al*. 2009; Isaacson and Scanziani 2011). In particular, the widespread inhibitory architecture described by Karnani *et al*. (2014) provides a plausible framework through which distributed inhibitory activity can coordinate communication across multiple cortical systems while preserving the overall organization of cortical networks.

Regional adjustments may arise indirectly through excitatory recruitment of local inhibitory circuits. Excitatory projections from auditory and motor cortices can engage VIP-mediated disinhibitory pathways in V1 (Iurilli *et al*. 2012; Fu et al. 2014), while sparse long-range GABAergic projections contribute to synchronizing distributed cortical activity (Tamamaki and Tomioka 2010; Melzer et al. 2012). Together, these mechanisms provide likely substrates for the distributed connectivity changes observed across inhibitory networks following early visual deprivation.

Although inhibitory functional networks appeared relatively stable at global and modular scales, nodal influence and modular analyses revealed subtle shifts in connector nodes (M2, AC) and bridging nodes (A, AM). These localized adjustments suggest that inhibitory circuits participate in sustaining modular organization when visual experience is absent and cannot guide the maturation of large-scale cortical structure. Inhibitory circuits are critical for preserving modular integrity and constraining excitatory spread (Isaacson and Scanziani 2011; Froemke 2015). Thus, rather than representing strict stability, we propose that inhibitory remodeling reflects targeted rebalancing of connectivity, maintaining local regulation within sensory areas while strengthening long-range couplings from associative cortices. This configuration illustrates the dual role of inhibitory networks as local stabilizers and global synchronizers, buffering excitatory perturbations and supporting cortical resilience under deprivation (Petrus *et al*. 2015).

Finally, inhibitory networks exhibited increased functional connectivity between frontal associative regions and medial visual cortices, consistent with the distributed pattern of connectivity reorganization observed across multiple functional systems. Similar fronto-occipital strengthening has been reported in early blind humans (Ortiz-Terán et al. 2017; Abboud and Cohen 2019), supporting the idea that inhibitory circuits contribute to coordinating long-range cortical communication under sensory deprivation. Together with the relative preservation of nodal influence and modular organization, these findings indicate that inhibitory cortical networks were modified primarily through selective redistribution of interregional communication rather than extensive topological remodeling. Such a reorganization is consistent with a stabilizing role for inhibitory circuits, allowing large-scale cortical integration to be maintained despite profound sensory deprivation.

## Pan-neuronal (hSyn) Population: Global Stability with Experience-Dependent Redistribution of Connectivity

In the pan-neuronal network, as previously reported (Laliberté and Boire 2026), early visual deprivation induced widespread redistribution of functional connectivity despite relative preservation of global network topology. The absence of significant changes in global network metrics despite extensive redistribution of individual connections is consistent with models of homeostatic structural plasticity, in which reorganization arises primarily through the reassignment of existing connections rather than the creation of new ones (Marder and Goaillard 2006; Yin and Yuan 2015; Makin and Krakauer 2023; Saccone *et al*. 2024). Because hSyn expression encompasses all neuronal classes (Nathanson et al. 2009), the resulting activity patterns likely approximate whole-cortex signals measured with BOLD fMRI or EEG, which similarly integrate activity across neuronal populations rather than isolating specific cell types. The preservation of global integration may further reflect metabolic and structural constraints, as cortical networks must balance wiring cost, energy expenditure, and computational efficiency (Karbowski 2014; Roberts et al. 2014; Herculano Houzel and Rothman 2022). Under sensory deprivation, large-scale remodeling therefore occurs within strict energetic limits, favoring redistribution of existing pathways over costly expansion.

Consistent with the nodal influence analyses, modular reorganization was characterized by a redistribution of influential regions from primary and lateral visual cortices toward medial higher visual and associative territories. At the network level, this shift was accompanied by partial fusion of previously distinct functional modules, reminiscent of the convergence between visual and language networks reported in blind humans (Hasson et al. 2016). Together, these observations suggest that sensory deprivation promotes broader multimodal integration while preserving overall cortical organization. Importantly, this convergent reorganization was consistently observed across functional connectivity, nodal influence, and modular analyses, reinforcing the interpretation that pan-neuronal cortical organization adapts through coordinated redistribution rather than global topological disruption.

In addition, homotopic interhemispheric connectivity was enhanced in blind mice, contrasting with the reduced homotopic interhemispheric connectivity of occipital visual cortices consistently reported in blind humans (Butt et al. 2015; Hou *et al*. 2017). This divergence likely reflects the denser callosal projections in rodents compared to primates (Ypma and Bullmore 2016), suggesting that the stronger bilateral scaffolding preserves or even amplifies interhemispheric coupling under visual deprivation.

These findings are consistent with the connectivity-constrained, experience-dependent hypothesis, which posits that functional connectivity emerges from the joint influence of anatomical architecture and sensory input (Saccone *et al*. 2024). Within this framework, the pan-neuronal network captures the synergistic interplay between excitatory and inhibitory populations, whereby extensive regional reorganization is accommodated without disrupting global cortical organization. Thus, the preservation of large-scale topology reflects coordinated reallocation of network roles that maintains efficient cortical communication despite profound sensory deprivation.

## Methodological Considerations and Limitations

Several methodological considerations should be acknowledged when interpreting the present findings. Because neuronal populations were compared using both transgenic and AAV-based expression strategies, differences in expression levels, signal-to-noise ratio, spatial coverage, and physiological effects associated with viral transduction may contribute to inter-cohort differences in functional connectivity metrics. To address this issue, spontaneous event dynamics were quantified across cohorts, revealing broadly comparable baseline activity profiles across promoter systems in sighted animals. In addition, all analyses were conducted primarily within the promoter, thereby limiting direct dependence on inter-cohort signal comparability. Nevertheless, because neuronal population and expression strategy were not independently manipulated, their respective contributions to the observed Promoter × Visual Status interactions cannot be fully dissociated.

An additional limitation is the relatively small and uneven sample sizes across experimental groups, particularly the hSyn enucleated cohort. Although the multivariate framework, permutation testing, and FDR correction were specifically implemented to reduce the risk of false-positive findings under these conditions, limited statistical power may have reduced sensitivity to detect more subtle effects, particularly at the level of global network metrics. Accordingly, negative findings, especially within the hSyn population, should be interpreted with appropriate caution.

We also did not directly assess astroglial or microglial activation following neonatal enucleation or viral vector administration. However, neonatal enucleation was performed at birth whereas imaging acquisitions began several weeks later (P70–P90), reducing the likelihood that acute inflammatory responses associated with optic nerve injury persisted during recordings. Similarly, viral vectors were administered intravenously rather than through direct cortical injections, thereby minimizing focal cortical inflammatory responses. Nonetheless, subtle long-term glial or inflammatory contributions to functional connectivity alterations cannot be completely excluded.

Finally, global signal regression was used to reduce widespread fluctuations associated with awake mesoscale imaging and improve the specificity of regional functional correlations. This preprocessing step was applied because awake mesoscale calcium imaging is particularly sensitive to global fluctuations arising from movement, respiration, photobleaching, LED drift, and optical inhomogeneity associated with transcranial imaging (Brier and Culver 2023). In addition, hemodynamic correction was performed prior to GSR to reduce vascular-related signal contamination. Although GSR improves regional specificity, it can alter covariance structure and introduce negative correlations. Connectivity patterns should therefore be interpreted within the methodological constraints associated with this preprocessing strategy.

## Conclusion

Our findings indicate that early blindness reshapes spontaneous cortical networks through both shared and neuronal identity-dependent mechanisms. Across neuronal populations, functional connectivity, nodal influence, and modular organization converged toward a common reorganization pattern characterized by reduced influence of primary and lateral visual areas together with enhanced integration of medial higher visual and associative cortices. Despite this shared organizational motif, neuronal populations differed markedly in how this reorganization was implemented: excitatory projection networks underwent the most extensive topological remodeling, inhibitory networks primarily redistributed communication while preserving large-scale network topology, and pan-neuronal networks maintained global organization through coordinated redistribution across complementary neuronal populations. These findings parallel evidence from early blind humans showing enhanced integration of occipital cortices within associative networks (Collignon *et al*. 2011) and are consistent with studies linking resting-state functional organization to underlying structural connectivity (Vincent *et al*. 2007; Honey *et al*. 2009). Together, our results identify neuronal identity as a key determinant of large-scale cortical plasticity and demonstrate that spontaneous cortical activity provides a powerful framework for understanding how sensory deprivation reshapes functional brain organization.

## Acknowledgements

We are grateful to the Animal Care Facilities staff, Nadia Desnoyers, Michel Demers, Christel Perron, and Vicky Renier-Tellier, for their support in animal care and maintenance. We also thank the *Service de consultation en statistique* of the Université du Québec à Trois-Rivières (UQTR), and especially Khadija Ouaqdi, for her expert statistical consultation and assistance with the analytical framework of this study.

