## Supplementary figures and images for "Excitatory and Inhibitory Networks Diverge Following Early Blindness"

### Supplementary Figure 1

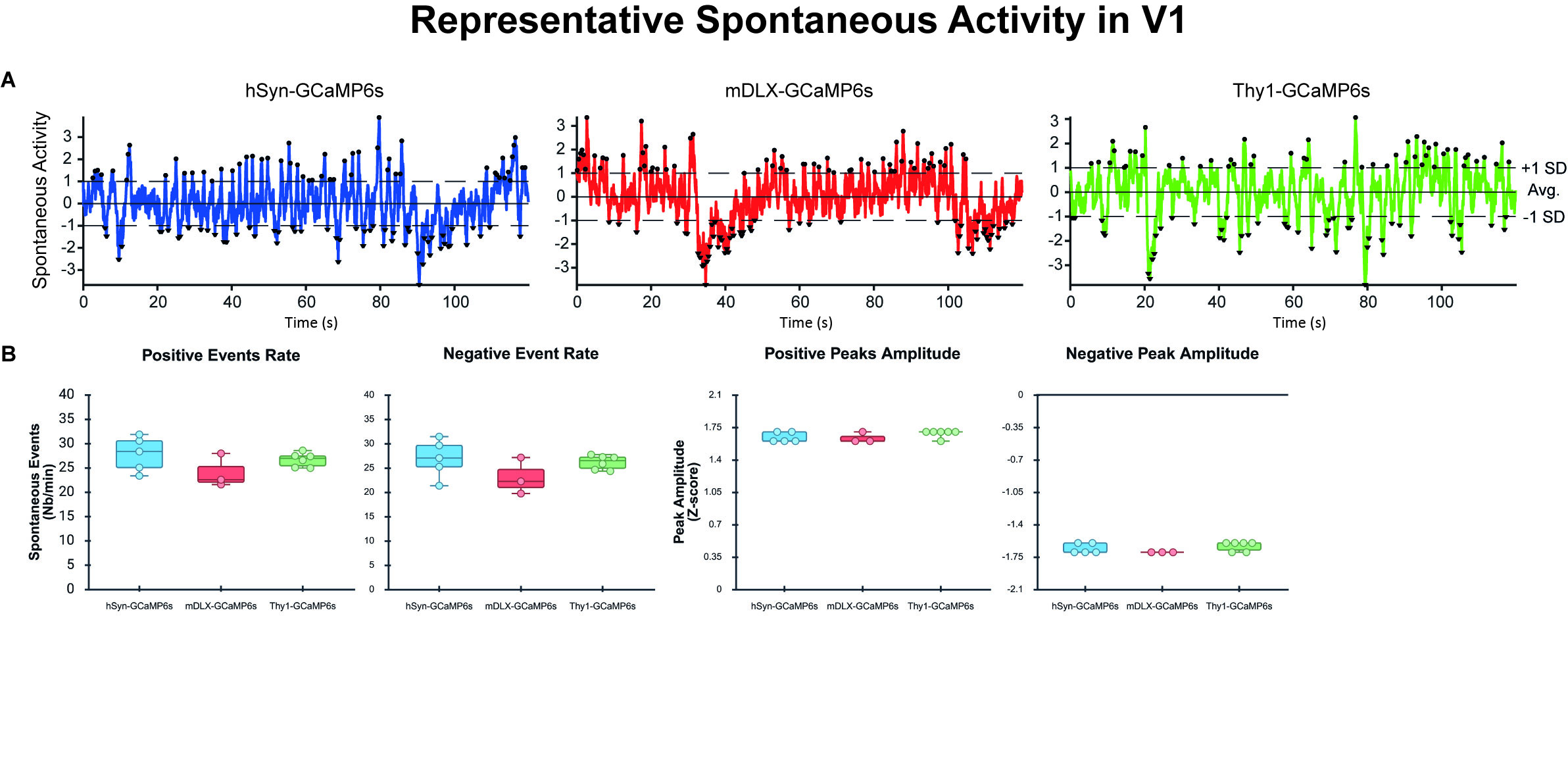
